# Spatial 3D genome organization controls the activity of bivalent chromatin during human neurogenesis

**DOI:** 10.1101/2024.08.01.606248

**Authors:** Sajad Hamid Ahanger, Chujing Zhang, Evan R. Semenza, Eugene Gil, Mitchel A. Cole, Li Wang, Arnold R. Kriegstein, Daniel A. Lim

**Author notes:** Equal contribution (Sajad Hamid Ahanger, Chujing Zhang).

## Abstract

The nuclear genome is spatially organized into a three-dimensional (3D) architecture by physical association of large chromosomal domains with subnuclear compartments including the nuclear lamina at the radial periphery and nuclear speckles within the nucleoplasm^1–5^. However, how spatial genome architecture regulates human brain development has been overlooked owing to technical limitations. Here, we generate high-resolution maps of genomic interactions with the lamina and speckles in cells of the neurogenic lineage isolated from midgestational human cortex, uncovering an intimate association between subnuclear genome compartmentalization, chromatin state and transcription. During cortical neurogenesis, spatial genome organization is extensively remodeled, relocating hundreds of neuronal genes from the lamina to speckles including key neurodevelopmental genes bivalent for H3K27me3 and H3K4me3. At the lamina, bivalent genes have exceptionally low expression, and relocation to speckles enhances resolution of bivalent chromatin to H3K4me3 and increases transcription >7-fold. We further demonstrate that proximity to the nuclear periphery – not the presence of H3K27me3 – is the dominant factor in maintaining the lowly expressed, poised state of bivalent genes embedded in the lamina. In addition to uncovering a critical role of subnuclear genome compartmentalization in neurogenic transcriptional regulation, our results establish a new paradigm in which knowing the spatial location of a gene is necessary to understanding its epigenomic regulation.

## Main

The three-dimensional (3D) organization of the human genome within the nucleus plays a key role in regulation of gene expression, which in turn controls cell differentiation and development ^6–8^. This 3D genome organization involves interaction of the genome with subnuclear structural compartments such as the nuclear lamina and nuclear speckles ^1,3^. The nuclear lamina lies underneath the inner nuclear membrane and is composed of lamin intermediate filament proteins ^9^. Roughly 30-40% of the human genome is anchored to the lamina through variably sized (10 kb–10 Mb) lamina-associated domains (LADs), and gene expression within LADs is generally low ^10–12^. Interior to the lamina, nuclear speckles are phase-separated, membrane-less compartments that contain a multitude of transcriptional regulators and RNA processing and splicing factors ^13–15^. Approximately, 10-15% of the human genome is organized around speckles, and genes within speckle-associated domains (SPADs) are more highly expressed ^10,16,17^. Thus, spatial genomic location with respect to subnuclear compartments is a key component of transcriptional control.

Previous studies have highlighted the role of LAD dynamics in different cell culture models, revealing a constitutive nature of LADs across these different cell lines as well as cell-type specific variability in genome-lamina interactions ^18,19^. In our previous work, we studied LAD architecture of the developing human, mouse, and macaque brains and showed that many LADs described as ‘constitutive’ in cell culture models are not observed in cells isolated from these tissues ^10^, highlighting the importance of studying cell lineages *in vivo*. SPADs have been studied in a handful of human cell lines, which revealed a largely conserved SPAD architecture across these cell lines ^16,20^. We have also profiled SPADs (along with LADs) in the developing human brain and found them to be enriched in genetic risk loci associated with schizophrenia ^10^. The organization of LADs and SPADs and how their dynamics contribute to lineage-specific gene regulation during the development of human brain remains unknown.

The control of gene expression during lineage-specification and cell differentiation also involves local chromatin modifications ^21^. Histone H3 lysine-4 trimethylation (H3K4me3) at gene promoters is associated with transcriptional activation, whereas histone H3-lysine-27 trimethylation (H3K27me3) marks repressed genes ^22^. The discovery of bivalent chromatin state, characterized by the local enrichment of both H3K4me3 and H3K27me3, added a new paradigm to developmental regulation of gene expression ^23,24^. Chromatin bivalency is thought to confer a lowly expressed but ready to be induced, often called as ‘poised’ transcriptional state to lineage-specifying genes in undifferentiated cells ^24–26^. Resolution of bivalency to either the H3K4me3- or H3K27me3-monovalent state is associated with gene activation or repression, respectively, during cell differentiation ^24–26^. However, it has been observed that the resolution of bivalent chromatin to the H3K4me3-monovalent state during differentiation does not always confer transcriptional induction ^25,27^. This indicates that impacts of bivalency on transcription are shaped by additional unknown factors. Whether 3D spatial genome organization and the bivalent chromatin state interact to regulate gene expression during development has not been investigated.

We recently developed Genome Organization with CUT&RUN technology (GO-CaRT), which enabled the mapping of LADs and SPADs with relatively small numbers (<100,000) of cells acutely isolated from the mouse, macaque, and human brains ^10^. Here we further develop this method for the study of specific cell types isolated from the human cortical neurogenic lineage, by first performing fluorescence-activated nuclei sorting (FANS) followed by GO-CaRT for mapping of LADs and SPADs. We addressed the methodological difficulty of using two antibody-based steps (FANS and GO-CaRT) by using fragment-antigen binding (Fab) to block the cell lineage specific antibody before performing the GO-CaRT to prevent spurious pA/G-MNase targeting. This method enabled us to study the organization and dynamics of LADs and SPADs during human prenatal cortical neurogenesis and the interplay between 3D spatial positioning in the nucleus and local chromatin modifications. Combining LaminB1 GO-CaRT with sequential chromatin immunoprecipitation (ChIP-reChIP) allowed us to study spatial 3D genome localization and chromatin signatures on the same DNA locus. Collectively, our results reveal a new mechanistic model in which subnuclear genome compartmentalization plays a dominant role in determining the overall transcriptional output of the bivalent genes important for neuronal development and cell differentiation.

## Results

### Global LAD architecture is extensively remodeled during human cortical neurogenesis

In the developing human cerebral cortex, radial glia (RG) – the major neural stem cell population – give rise to proliferative intermediate progenitor cells (IPC) that then produce post-mitotic excitatory neurons (EN) ^28,29^. Using fluorescence-activated nuclei sorting (FANS) with antibodies to PAX6, EOMES, and SATB2, we isolated nuclei of RG (PAX6+), IPC (PAX6+, EOMES+), and EN (SATB2+) cell populations from primary cortical brain tissue of gestational week (GW) 20 (**Fig. 1a, Extended Data Fig. 1a**). RNA-seq analysis performed on nuclei isolated by FANS confirmed the separation of these cell types and their identity (**Extended Data Fig. 1b**).

**Figure 1.**
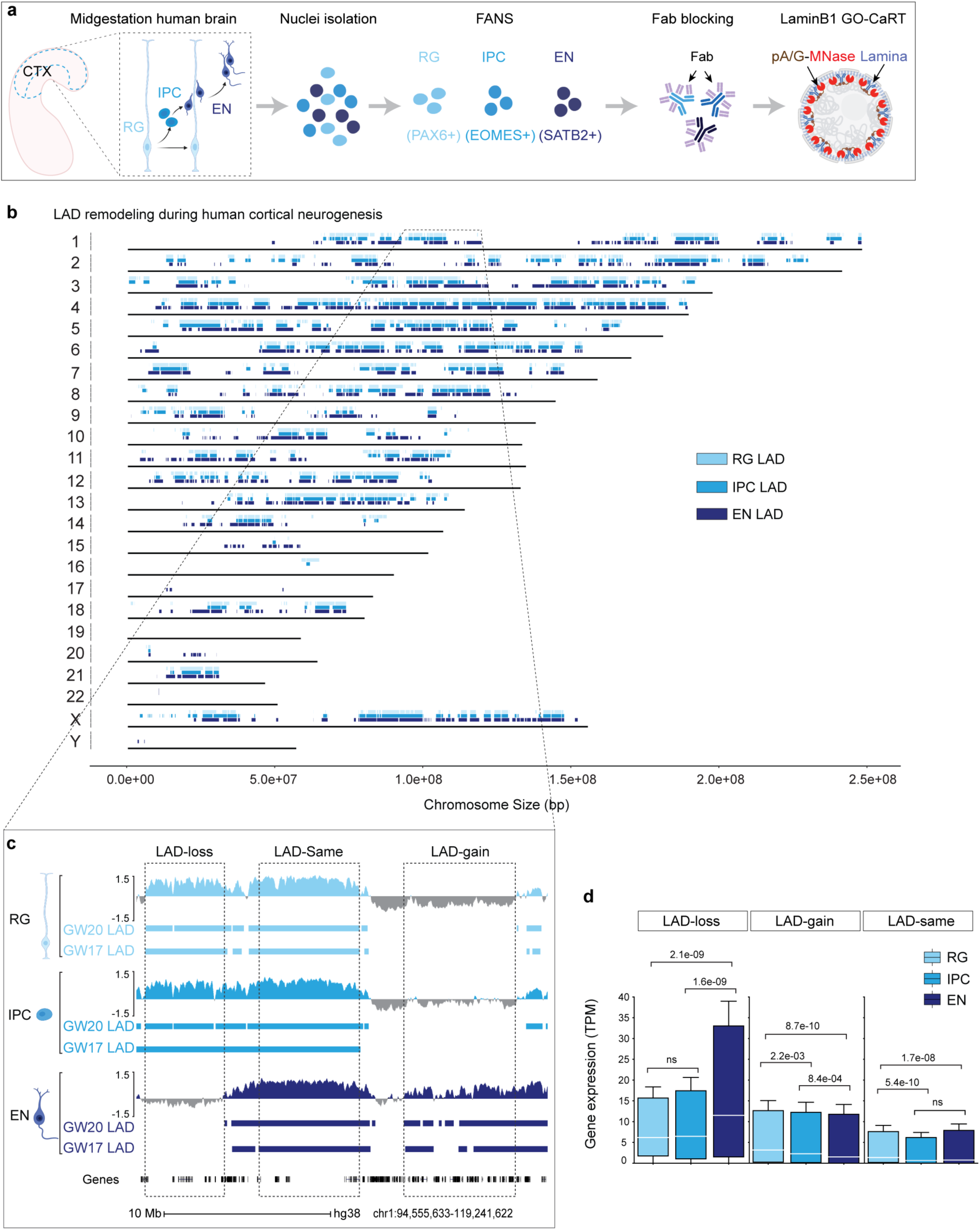
LAD architecture is extensively remodeled during human cortical neurogenesis. **a,** Schematic of workflow for FANS-based isolation of RG, IPC and EN, Fab blocking of antibodies used in FANS and downstream LaminB1 GO-CaRT analyses. **b,** Distribution of LADs across all the human chromosomes at GW17 and GW20. LADs that were identified in both GW17 and GW20 samples are indicated by rectangles in light blue for RG, medium light blue for IPC and dark blue for EN. **c,** Representative LaminB1 GO-CaRT tracks in RG, IPC and EN at GW20. The Y-axis depicts log2 ratio of LaminB1 over IgG. LAD calls at GW20 and GW17 are depicted below tracks by rectangles. Dashed boxes show example genomic regions where LADs are lost (LAD-loss), gained (LAD-gain) or remain unchanged (LAD-same) during RG to EN differentiation. **d,** Box plot showing average gene expression (in transcripts per million, TPM) in LAD-loss (n = 1240), LAD-gain (n = 3241) and LAD-same regions (n = 3038) *n*, number of genes; *P* values determined by Wilcoxon rank sum test. Boxes show the range from lower quartiles (25th percentile) to upper quartiles (75th percentile), with the median line (50th percentile); whiskers extend 1.5 times the interquartile range from the bounds of the box. Related data in Extended Data Fig. 1, 2 and 3.

In GO-CaRT – a chromatin profiling technology based on CUT&RUN – antibodies that bind proteins in specific nuclear compartments are used to target protein A/G-fused micrococcal nuclease (pA/G-MNase), resulting in the cutting and release of compartment-associated genomic DNA which is then identified with next generation sequencing ^10^. We used monovalent fragment antigen-binding (Fab) fragments, which lack the fragment crystallizable (Fc) domain to which pA/G binds, to “block” antibodies used in FANS prior to the GO-CaRT protocol. Fab fragments were highly efficient at blocking undesired antibody interactions without interfering with downstream GO-CaRT assays (**Extended Data Fig. 2a-f**). After Fab blocking, we performed LaminB1 GO-CaRT on FANS-isolated nuclei (∼25,000) from GW17 and GW20 cortex, and the LaminB1 profiles from RG, IPC and EN populations were highly similar between GW17 and GW20 (**Fig. 1a, Extended Data Fig. 3a**), indicating that these developmental time points are reasonable biological replicates for studies of LAD architecture.

To identify LAD architectural changes related to human cortical neurogenesis, we compared the LAD profiles of RG, IPC and EN (**Fig. 1b)**. LAD coverage in cells across this neurogenic lineage was 36-38% of the genome, and these lamina-associated regions exhibited genomic features typical of LADs including lower gene density and expression (**Extended Data Fig. 3b-d**). During the differentiation of RG to EN, the global LAD architecture was extensively remodeled (**Fig. 1b, c, Extended Data Fig. 3e**). Specifically, 322 Mb (10.4% of the whole genome) moved from the nuclear interior to the lamina (LAD-gain), 253 Mb (8.2% of the whole genome) were released from the lamina to the interior space of the nucleus (LAD-loss) and 778 Mb (25.2% of the whole genome) remained unchanged (LAD-same).

The lamina is a nuclear compartment that is generally repressive to transcription^10,12^. Across the three cell types of the neurogenic lineage, genes in LAD-same regions exhibited significantly low levels of RNA expression (**Fig. 1d**). In contrast, genes in LAD-loss and LAD-gain regions had significantly increased or decreased gene expression, respectively, in EN as compared to RG (**Fig. 1d**). Of note, LAD-gain regions (3241 genes/322 Mb) had approximately twice the gene density as that of LAD-loss regions (1260 genes/253 Mb) (**Extended Data Fig. 3f**), indicating that during differentiation into postmitotic neurons, a greater number of genes are moved into LADs than released from the lamina. Thus, during human cortical neurogenesis, LAD architecture is dynamic and involves large-scale movement of genomic regions from and to the nuclear lamina.

### SPAD architecture of postmitotic excitatory neurons is distinct from that of proliferative neural progenitors

The 3D genome organization around nuclear speckles has been analyzed in only a few human cell lines and bulk human cortical tissue ^10,17,20^. To examine the SPAD architecture of human neurogenic cell lineage, we performed GO-CaRT using an antibody against SON (a nuclear speckle scaffold protein) in RG, IPC and EN at GW17 (**Fig. 2a**). In RG, IPC and EN, the SPAD coverage was between 15-25% of the genome. Across the neurogenic lineage, each chromosome was organized into largely mutually exclusive SPADs and LADs with intervening non-LAD/non-SPAD regions (**Fig. 2b**). SPAD genes were expressed at higher levels than those in LADs or non-LAD/non-SPAD regions (**Fig. 2c)**. Furthermore, genes within SPADs were longer, contained more exons, and the transcripts had greater isoform diversity (**Fig. 2c, Extended Data Fig. 4a**). Thus, across the neurodevelopmental lineage, SPAD genes exhibit increased expression and splicing activity.

**Figure 2.**
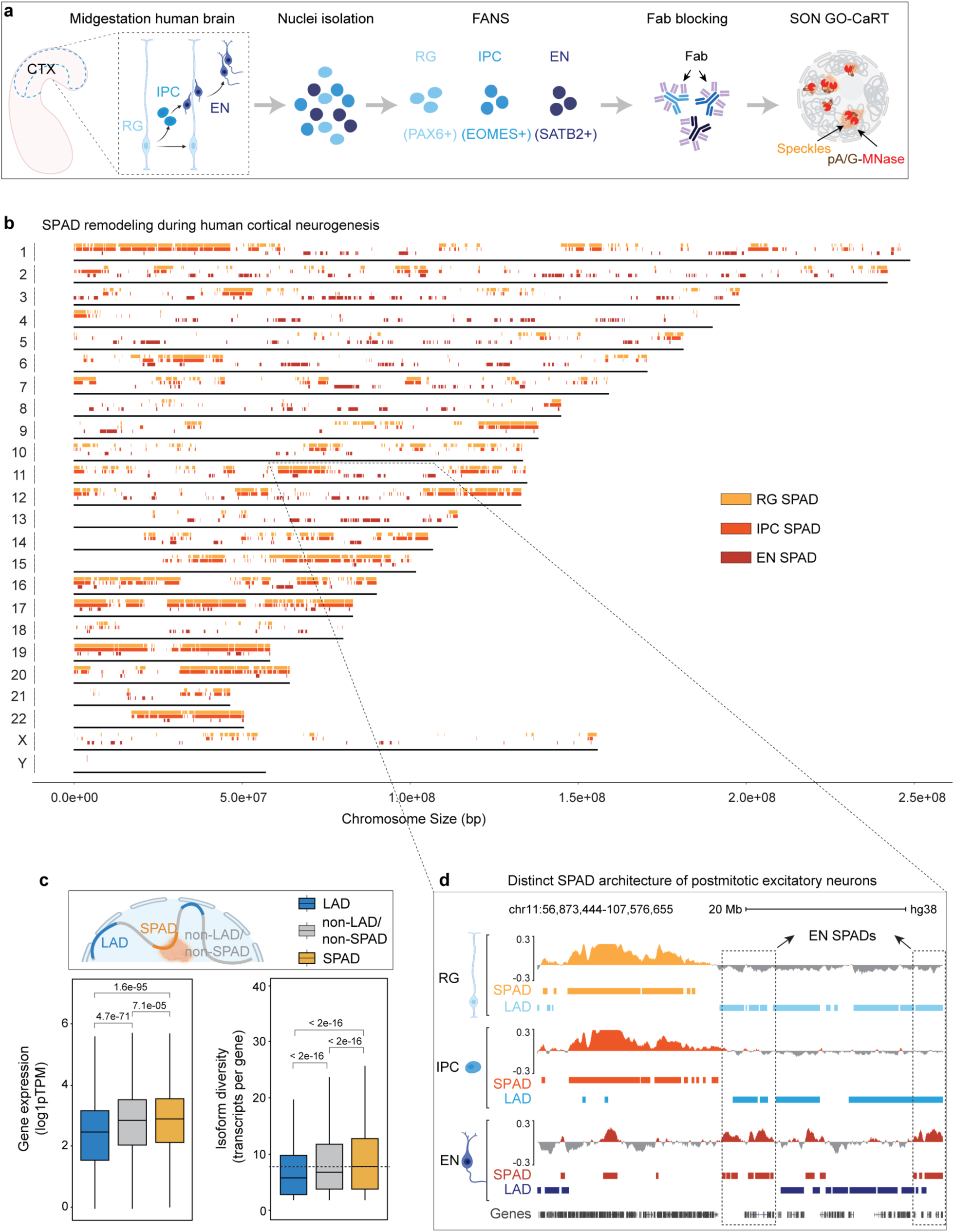
The SPAD architecture of postmitotic EN is distinct from that of proliferative RG and IPC. **a,** Schematic of workflow for FANS-based isolation of RG, IPC and EN, Fab blocking of antibodies used in FANS and downstream SON GO-CaRT analyses. **b,** Distribution of SPADs across all the human chromosomes at GW17. SPADs are indicated by rectangles of light orange for RG, medium light orange for IPC and dark orange for EN. **c,** Box plot showing average gene expression and isoform diversity in LADs (n = 6367), non-LADs/non-SPADs (n =14962) and SPADs (n = 14879) across the neurogenic lineage. Non-LADs/non-SPADs are genomic regions that neither associate with lamina nor speckles. *n*, number of genes; *P* values determined by Wilcoxon rank sum test. Boxes show the range from lower quartiles (25th percentile) to upper quartiles (75th percentile), with the median line (50th percentile); whiskers extend 1.5 times the interquartile range from the bounds of the box. **d,** Representative SON GO-CaRT tracks in RG, IPC and EN showing differences in SPAD architecture between RG/IPC and EN (dashed boxes). SPAD (shades of orange) and LAD calls (shades of blue) are depicted below the tracks for comparison. Related data in Extended Data Fig. 4.

While SPAD profiles were highly similar between RG and IPC (Jaccard = 0.59), the SPAD architecture of EN was more distinct (Jaccard: 0.04-0.06, **Fig. 2b, d, Extended Data Fig. 4b**). Gene ontology (GO) analyses revealed that in RG and IPC, SPADs were similarly enriched in genes related to nuclear processes (transcription, replication, RNA processing, chromatin organization etc.), metabolic processes, cytoskeleton organization and cell cycle regulation (**Extended Data Fig. 4c**). In contrast, EN SPADs were predominantly enriched in genes involved in neuronal pathways including neuron projection, axon guidance and synapse function (**Extended Data Fig. 4c**). Thus, in the developing human cortex, the SPAD architecture of postmitotic excitatory neurons is distinct from that of proliferative neural progenitors (RG, IPC).

### Neuronal genes detach from the lamina and relocate to speckles during cortical neurogenesis

After release from the nuclear lamina, certain genomic regions can become associated with nuclear speckles. During RG to EN differentiation, 1240 genes were released from the nuclear lamina. In these LAD-loss regions, GO analyses revealed an enrichment of genes involved in neuronal processes including axonogenesis, cell-cell adhesion, neuron projection guidance, synapse assembly and nervous system development (**Fig. 3a**). Included in this set of genes were synaptic genes such as *ANKS1B, NRG3, SYT1, SLITRK1,* and *LRRC7* and transcription factors with well-known roles in cortical development such as *MEF2C* and *SATB2*. Genes (n= 3241) that gained lamina association during RG to EN differentiation were not significantly enriched in any specific cellular pathways by GO analysis, but we note that key transcription factors involved in RG function (*e.g., SOX2, HOPX*), multipotent state (*e.g., NANOG)*, neural precursor state (*e.g., ASCL1*) non-excitatory neuron lineages (*e.g., NKX2-1, NKX2-2,* and *GSX2*), and non-neural fates (*e.g., MYOD1* and *ZEB1*) are in LAD-gain regions (**Extended Data Fig. 5a, b**). Thus, LAD remodeling during RG to EN differentiation involves sequestration of non-neural lineage genes to the lamina and the release of genes important for neuronal differentiation and function.

**Figure 3.**
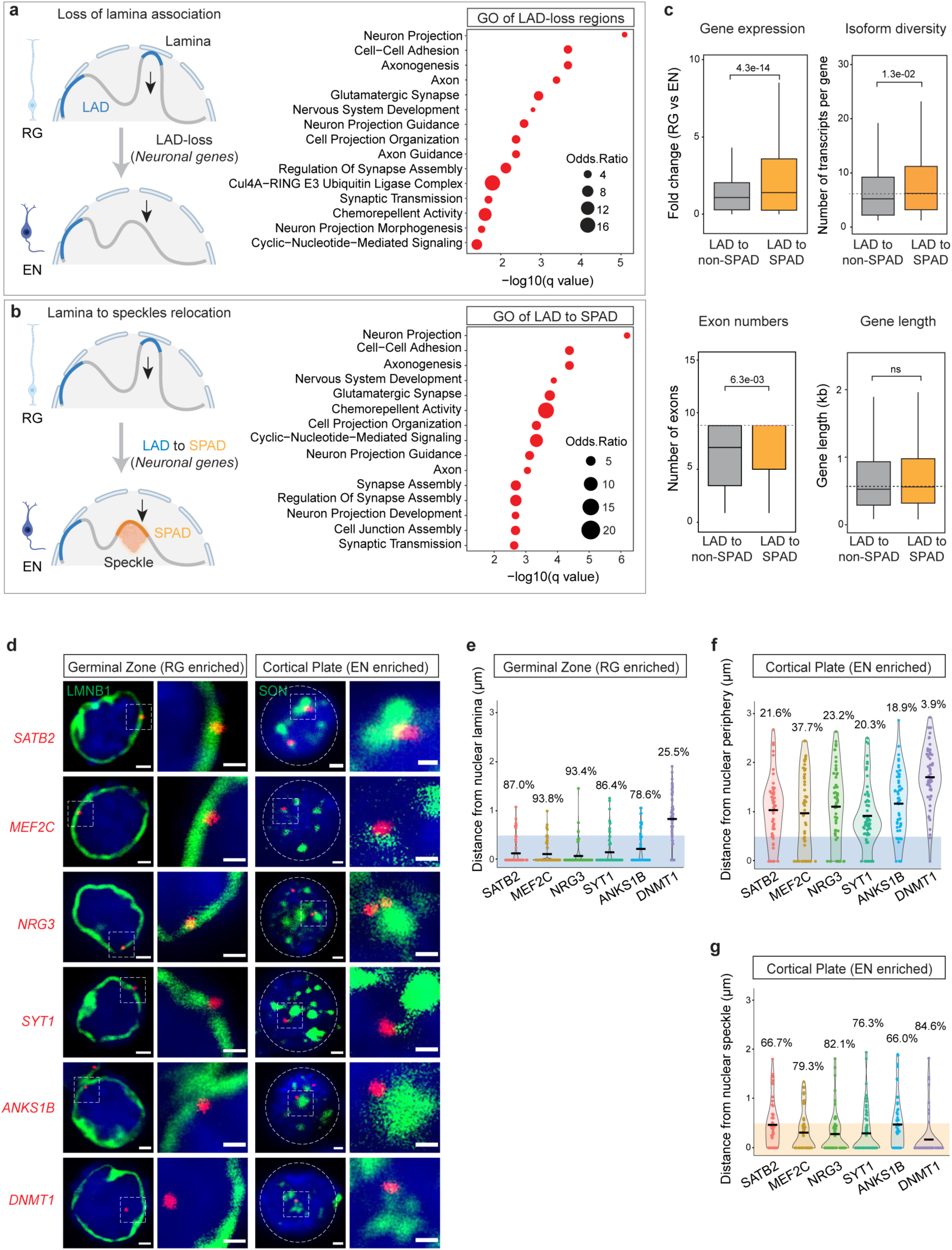
Neuronal genes move from lamina to speckles during cortical neurogenesis. **a,** A cartoon illustrating LAD-loss during RG to EN differentiation and GO analysis of these regions. Top 15 enriched GO terms (biological processes) are shown. GO terms were sorted based on their significance (−log10(*q* value)); the size of the bubble represents the Odds Ratio for each term. **b,** A cartoon illustrating LAD to SPAD switch during RG to EN differentiation and GO analysis of these regions. Top 15 enriched GO terms (biological processes) are shown. **c,** Box plot showing average gene expression, isoform diversity, exon number and gene length in regions that switch from LAD to non-SPAD (n = 544) and LAD to SPAD (n = 696) during RG to EN differentiation. *n*, number of genes; *P* values determined by Wilcoxon rank sum test. Boxes show the range from lower quartiles (25th percentile) to upper quartiles (75th percentile), with the median line (50th percentile); whiskers extend 1.5 times the interquartile range from the bounds of the box. **d,** Representative micrographs of DNA-FISH of indicated gene loci (in red) with LaminB1 and SON immunostaining (in green) in cells of the germinal zone (GZ) and cortical plate (CP) at GW17. Scale bar, 2 μm; higher magnification, 0.5 μm; DAPI in blue. **e, f, g** Quantifications of DNA-FISH loci shown in D. Percentages indicate loci within 0.5 μm from the nuclear lamina (GZ), nuclear periphery (CP) or nuclear speckles (CP). For each locus, 50–60 nuclei were quantified. Related data in Extended Data Fig. 5 and 6.

Among the 1240 genes that were released from the nuclear lamina during RG to EN differentiation, ∼56% (696 genes) relocated to nuclear speckles. GO analyses revealed this set of SPAD genes to be enriched in processes related to synaptic function, cell-adhesion, and axon guidance (**Fig. 3b**). In contrast, the set of genes (544) that relocated to non-LAD/non-SPAD regions were not significantly enriched in any specific cellular pathway. Furthermore, in comparison to non-LAD/non-SPAD regions, genes that relocated to SPADs were expressed at higher levels, contained more exons, and had greater isoform diversity (**Fig. 3c**). These data suggest that nuclear speckles during cortical neurogenesis act as a hub for neuronal gene expression and splicing, particularly of genes involved in synapse function.

### DNA-FISH confirms the movement of neuronal genes from the lamina to speckles

Using DNA fluorescence *in situ* hybridization (FISH) combined with immunocytochemistry (ICC), we analyzed the nuclear localization of five neuronal gene loci – *SATB2, MEF2C, ANKS1B, NRG3,* and *SYT1 –* that were repositioned from lamina/LAD in RG to speckles/SPAD in EN (**Extended Data Fig. 6a**). For comparison, we also analyzed the nuclear localization of *DNMT1*, a non-LAD gene in both RG and EN. (**Extended Data Fig. 6a**). DNA-FISH was performed in cells dissociated from the germinal zone (which is enriched in RG) and cortical plate (which is enriched in neurons) from GW17 brain (**Extended Data Fig. 6b**). The cells from the germinal zone were stained for LaminB1 (to mark nuclear lamina) and those from the cortical plate were stained with SON (to mark nuclear speckles). Quantitative analysis of DNA-FISH data showed that *SATB2, MEF2C, ANKS1B, NRG3, and SYT1* loci were preferentially located close to the LaminB1+ nuclear periphery in cells of the germinal zone (**Fig. 3d, e**). In the cortical plate, these gene loci were distant to the nuclear periphery and instead localized close to SON+ nuclear speckles (**Fig. 3d, f, g**). The non-LAD *DNMT1* locus was localized away from the nuclear periphery and close to the nuclear speckles both in the cells of the germinal zone and the cortical plate (**Fig. 3D-G**). These results validate GO-CaRT data and the movement of neuronal genes from lamina to speckles during RG to EN differentiation.

### Lamina association contributes to the lowly expressed, ‘poised’ state of bivalent genes

The local presence of both H3K4me3 and H3K27me3 – known as bivalent chromatin state – is a key feature of many developmentally regulated gene promoters ^24,25^. While genes monovalent for H3K4me3 or H3K27me3 generally correspond to active or repressed gene expression, respectively, bivalent genes are “poised” at variably intermediate levels of gene expression ^24,30^. To examine whether localization at the lamina contributes to the lowly expressed, ‘poised’ state of bivalent genes, we performed CUT&RUN ^31^ for H3K4me3 and H3K27me3 in RG, IPC and EN and identified bivalent genes containing peaks for both of these marks. Because the bivalent chromatin state is a key feature of stem cells ^24,30^, we focused our analyses on RG. In RG, across the whole genome, 1759 genes were bivalent for H3K4me3-H3K27me3 out of which 272 were in LADs. Bivalent genes in RG LADs were enriched in various GO terms related to neuron differentiation and function (**Extended Data Fig. 7a**). In contrast, genes monovalent for H3K4me3 (n= 948) and H3K27me3 (n=243) in RG LADs were enriched in GO terms unrelated to neuronal differentiation (**Extended Data Fig. 7b, c**). Thus, H3K4me3-H3K27me3 bivalent genes in RG LADs are enriched in neurodevelopmental function.

In RG, genes monovalent for H3K4me3 were expressed 4.2-fold higher than those monovalent for H3K27me3, and bivalent genes were expressed at intermediate levels (**Fig. 4a**). Importantly, bivalent genes were expressed at significantly lower levels in LADs as compared to those in non-LADs and those across the whole genome (**Fig. 4a**). For H3K27me3-monovalent genes, location within LADs and non-LADs did not correspond to major differences in their low levels of expression (**Fig. 4a**). Genes monovalent for H3K4me3 were highly expressed, with significantly lower levels in LADs (**Fig. 4a**). Thus, the lower expression level of bivalent genes corresponds to LAD location, indicating that this spatial aspect of genome organization contributes to their lowly expressed, ‘poised’ state.

**Figure 4.**
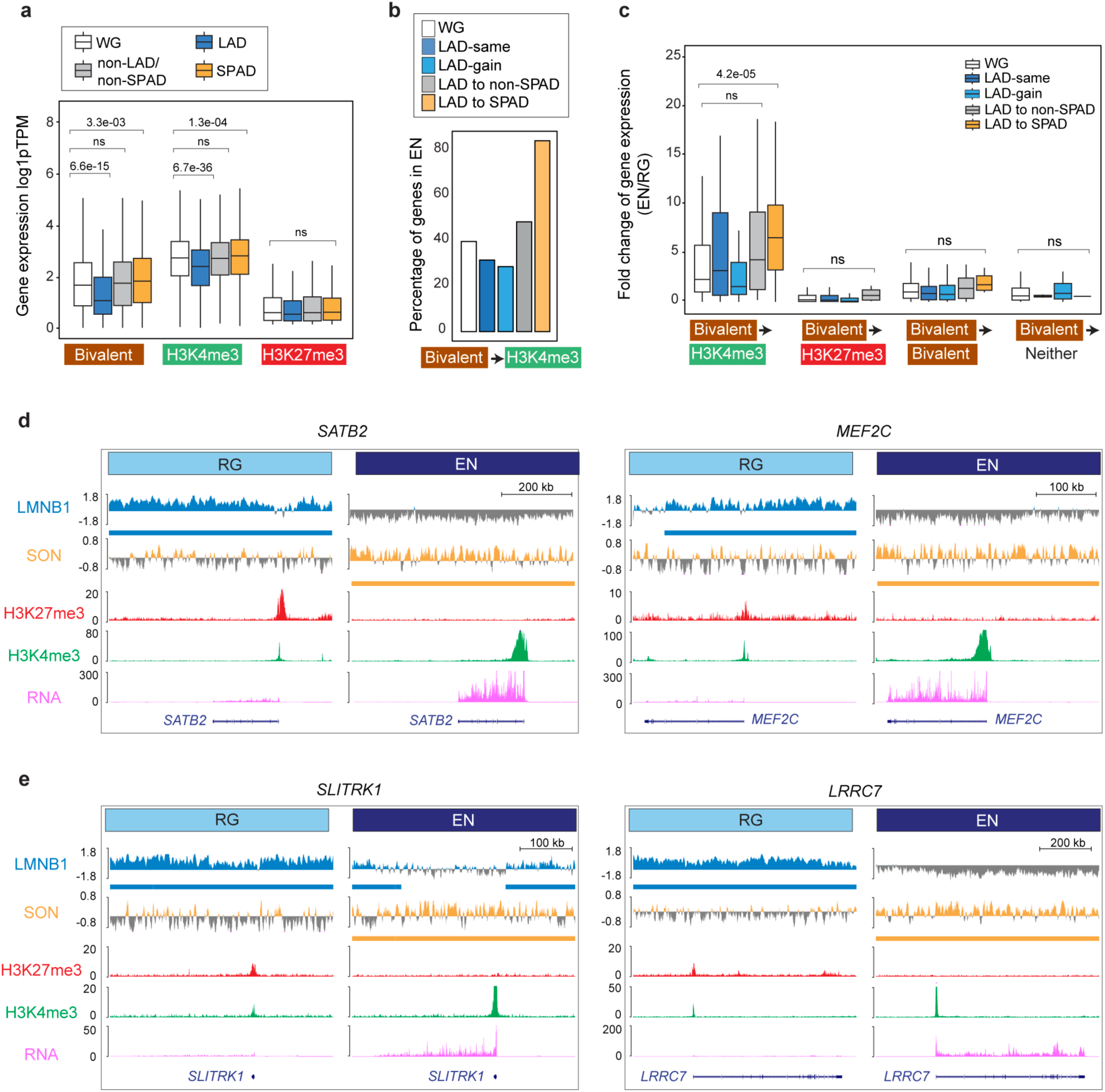
LAD and SPAD dynamics regulate the bivalent chromatin state. **a,** Box plot showing average gene expression of RG bivalent (n = 1873), H3K4me3-monovalent (n = 5014), and H3K27me3-monovalent (n = 582) genes in LADs, SPADs, non-LAD/non-SPADs and the whole genome (WG) in RG. *n*, number of genes across the WG; *P* values determined by Wilcoxon rank sum test. Boxes show the range from lower quartiles (25th percentile) to upper quartiles (75th percentile), with the median line (50th percentile); whiskers extend 1.5 times the interquartile range from the bounds of the box. **b,** Percentage of RG bivalent genes that resolve to the H3K4me3-monovalent state in EN based on LAD to SPAD dynamics. **c,** Fold-change of expression of bivalent genes that resolve to the H3K4me3-monovalent (n = 705), H3K27me3-monovalent (n = 207), remain bivalent (n = 802), or lose both marks (n = 43) during RG to EN differentiation based on LAD to SPAD dynamics. *n*, number of genes across the WG; *P* values determined by Wilcoxon rank sum test. Boxes show the range from lower quartiles (25th percentile) to upper quartiles (75th percentile), with the median line (50th percentile); whiskers extend 1.5 times the interquartile range from the bounds of the box. **d, e,** LaminB1 GO-CaRT, SON GO-CaRT, H3K27me3, H3K4me3 (CUT&RUN) and RNA-seq tracks at some example RG LAD bivalent genes that relocate to SPADs in EN and resolve to the H3K4me3-monovalent with strong transcriptional upregulation. These genes include key transcriptional factors, *SATB2*, *MEF2C* (D) and synapse genes, *SLITRK1* and *LRRC7* (E). Related data in Extended Data Fig. 7.

### LAD regions that relocate to speckles exhibit enhanced resolution of bivalent chromatin to active state

The “resolution” of bivalent genes to the H3K4me3-monovalent state corresponds to an increase in gene expression, particularly during cell differentiation. In RG, out of the 1759 bivalent genes, ∼40% resolved to monovalent H3K4me3 in EN, corresponding to an increase (2.3-fold) in gene expression (**Fig. 4b, c**). In both LAD-same and LAD-gain regions, fewer bivalent genes (28-30%) were resolved to H3K4me3-monovalent with a moderate increase in gene expression (**Fig. 4b, c**). In the subset of LAD-loss regions that relocated to non-LAD/non-SPAD regions, ∼48% of RG bivalent genes became monovalent for H3K4me3 and exhibited a greater increase in gene expression (4.3-fold) in EN as compared to RG (**Fig. 4b, c)**. In the subset of LAD-loss regions that relocated to EN SPADs, an overwhelming majority (83%) of the RG bivalent genes were resolved to the H3K4me3-monovalent state corresponding to further increase (7.1-fold) in gene expression (**Fig. 4b, c**). Examples of RG bivalent genes that relocated to EN SPADs included the key neurodevelopmental transcription factors *SATB2* and *MEF2C* and synaptic genes *SLITRK1* and *LRRC7* (**Fig. 4d, e**). Thus, the movement of genomic regions from the lamina to nuclear speckles during human cortical neurogenesis strongly corresponds to the resolution of resident bivalent genes to the H3K4me3-monovalent state and increased transcriptional output.

### LaminB1 GO-CaRT.ChIP-reChIP confirms the bivalent chromatin state of LAD genes

Parallel CUT&RUN (or similar) assays for H3K4me3 and H3K27me3 are unable to unequivocally establish the presence of both these marks on the same chromatin fragment. To further examine the bivalent chromatin state of LAD genes, we conducted LaminB1 GO-CaRT in neural precursor cells (NPCs) derived from human induced pluripotent stem cells (hiPSC, **Fig. 5a**). The LaminB1 GO-CaRT profile of iPSC-derived NPCs was highly similar to that of the RG (**Fig. 5b**, Spearman = 0.87). We next performed sequential chromatin immunoprecipitation (ChIP-reChIP), in which LaminB1 GO-CaRT-released chromatin was used as input for H3K27me3 ChIP, and the eluted chromatin from this first ChIP was subsequently used as input for H3K4me3 ChIP (LaminB1 GO-CaRT.ChIP-reChIP, **Fig. 5c, Extended Data Fig. 8a, b**). We also carried out ChIP-reChIP on LaminB1 GO-CaRT-released chromatin with the order of ChIP reversed (*i.e.,* H3K4me3 followed by H3K27me3). Of the LAD genes called “bivalent” by parallel H3K4me3 and H3K27me3 CUT&RUN analysis, LaminB1 GO-CaRT.ChIP-reChIP analysis confirmed ∼88% (338 out of 384) of them. In particular, transcription factor genes *SATB2* and *MEF2C* and synapse genes *SLITRK1* and *LRRC7* were confirmed to be bivalent by LaminB1 GO-CaRT.ChIP-reChIP (**Fig. 5d**). In contrast, *COL11A1* and *TFAP2D/B,* which localize to LADs in NPCs but are monovalent for H3K4me3 and H3K27me3 respectively, were not enriched by LaminB1 GO-CaRT.ChIP-reChIP (**Fig. 5d**), demonstrating the specificity of sequential ChIP on LaminB1-enriched chromatin. These results confirm that LADs contain bivalent chromatin at genes important for neuronal differentiation.

**Figure 5.**
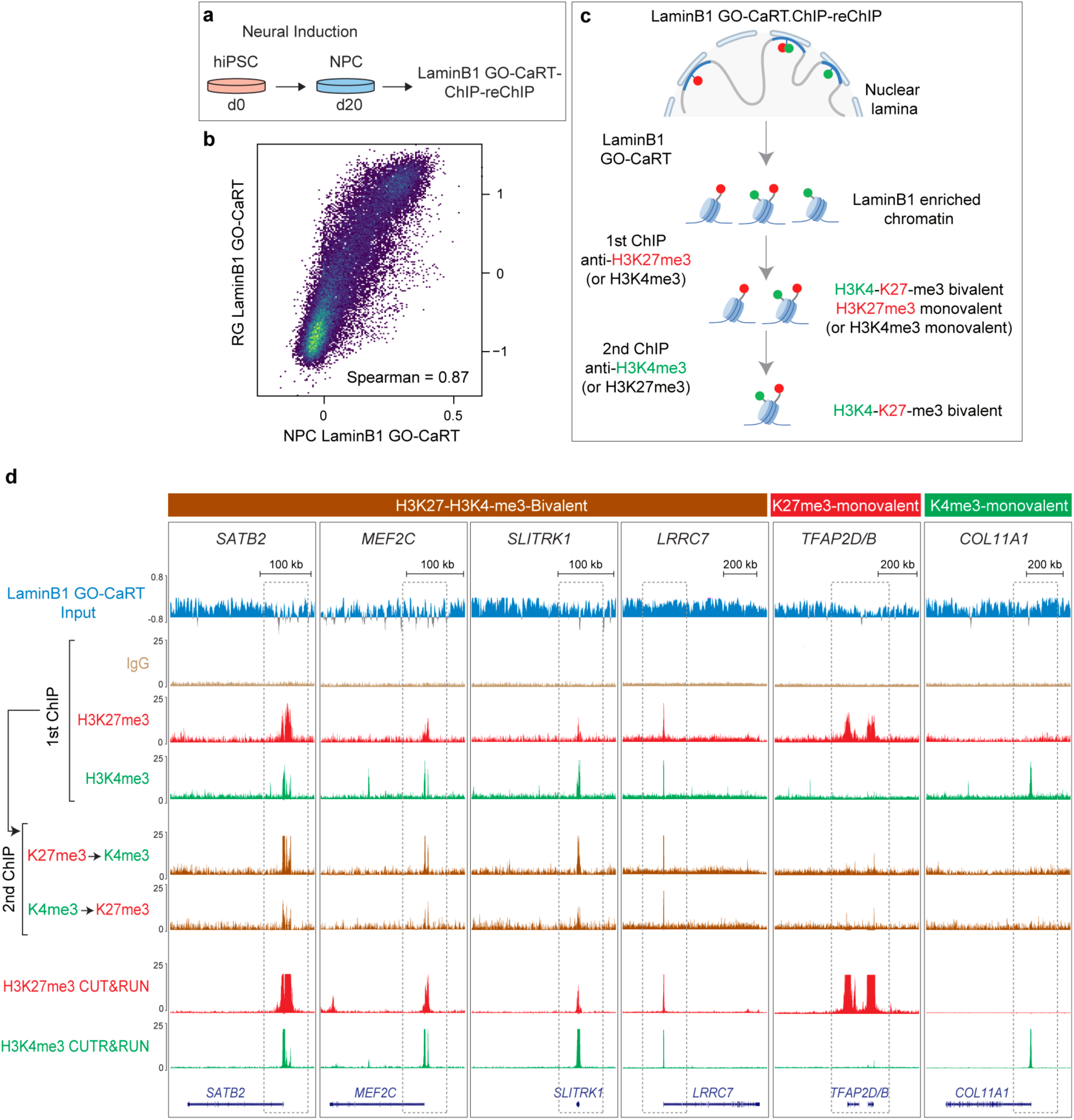
LaminB1 GO-CaRT.ChIP-reChIP confirms the bivalent chromatin state of LAD genes. **a,** Schematic of neural induction of human iPSCs and LaminB1 GO-CaRT-sequential ChIP (ChIP-reChIP) on iPSC-derived NPCs. **b,** Genome-wide scatter plot showing Spearman correlation between LaminB1 GO-CaRT in iPSC-derived NPCs and LaminB1 GO-CaRT in RG isolated from GW17 cortex. **c,** Schematic of LaminB1 GO-CaRT.ChIP-reChIP, in which LaminB1 GO-CaRT-released chromatin is used as input for a sequential chromatin immunoprecipitation (ChIP-reChIP), first with an antibody against H3K27me3 followed by a second ChIP with an antibody against H3K4me3. This ChIP-reChIP is also carried out with the order of ChIP reversed (*i.e.,* H3K4me3 ChIP followed by H3K27me3 ChIP). **d,** LaminB1 GO-CaRT.ChIP-reChIP profiles at key neuronal LAD genes, *SATB2*, *MEF2C*, *SLITRK1* and *LRRC7*. LAD genes monovalent for H3K27me3 (*TFAP2D/B*) and H3K4me3 (*COL11A1*) are not enriched by LaminB1 GO-CaRT.ChIP-reChIP. The following tracks are shown: LaminB1 GO-CaRT input, first ChIP with H3K27me3 and H3K4me3, H3K4me3 sequential ChIP on H3K27me3 first ChIP and H3K27me3 sequential ChIP on H3K4me3 first ChIP. IgG is a non-specific negative control for ChIP. Parallel H3K27me3 and H3K4me3 CUT&RUN tracks are also shown for comparison. Related data in Extended Data Fig. 8.

### LAD remodeling occurs early and regulates the activity of bivalent chromatin during *in vitro* neurogenesis

To further study the relationship between lamina-association and the bivalent chromatin state, we generated profiles of LADs, H3K4me3 and H3K27me3 during the synchronous differentiation of hiPSC-derived NPCs into cortical neurons (**Fig. 6a**) ^32^. In both NPCs and NPC-derived neurons (iN), LAD coverage was ∼34% of the genome. During neuronal differentiation (1 week), 173 Mb of the genome dissociated from the lamina (LAD-loss, 5.6% of the genome), 269 Mb gained lamina association (LAD-gain, 8.7% of the genome) and 781 Mb remained unchanged (LAD-same, 25.3% of the genome) (**Fig. 6b**). Genes in LAD-loss regions were enriched in GO terms related to neuronal processes including neuron projection, synapse assembly and axon guidance (**Extended Data Fig. 9a**). Thus, LAD remodeling occurs early during neuronal differentiation and releases genes important for neuronal function.

**Figure 6.**
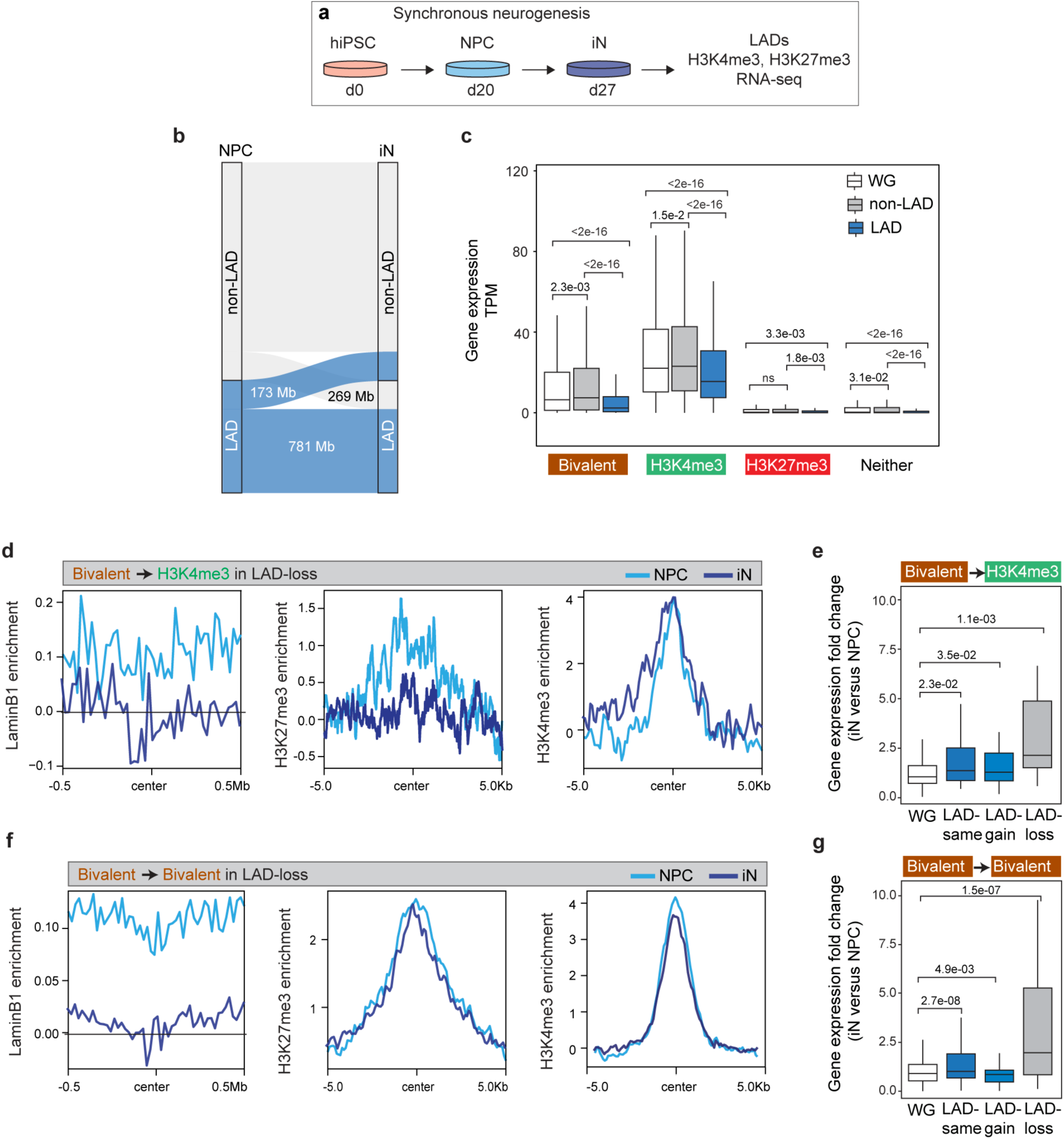
LAD remodeling occurs early and regulates the activity of bivalent chromatin during *in vitro* neurogenesis. **a,** Schematic of synchronous neuronal differentiation of iPSC-derived NPCs and profiling of LADs, histone marks and RNA-seq in NPC and induced neurons (iN). **b,** Alluvial plot depicting LAD changes (LAD-loss and LAD-gain) during the differentiation of NPC to iN. **c,** Box plot showing average gene expression of bivalent (n = 1024), H3K4me3-monovalent (n = 3522), H3K27me3-monovalent (n = 407) and genes containing neither mark in LADs, non-LADs and the WG in NPCs. *n*, number of genes across the whole genome; *P* values determined by Wilcoxon rank sum test. Boxes show the range from lower quartiles (25th percentile) to upper quartiles (75th percentile), with the median line (50th percentile); whiskers extend 1.5 times the interquartile range from the bounds of the box. **d,** Normalized LaminB1, H3K27me3 and H3K4me3 levels at bivalent genes that resolve to the H3K4me3-monovalent state in LAD-loss regions during NPC to iN differentiation. **e,** Fold-change of expression of bivalent genes that resolve to the H3K4me3-monovalent state in LAD-loss regions during NPC to iN differentiation. *P* values determined by Wilcoxon rank sum test. **f,** Normalized LaminB1, H3K27me3 and H3K4me3 levels at bivalent genes that retain bivalency in LAD-loss regions during NPC to iN differentiation. **g,** Fold-change of expression of bivalent genes that retain bivalency in LAD-loss regions during NPC to iN differentiation. *P* values determined by Wilcoxon rank sum test. Related data in Extended Data Fig. 9.

Similar to our findings in primary cells, bivalent genes in NPCs were expressed at significantly lower levels in LADs as compared to non-LADs or the whole genome (**Fig. 6c**). Genes monovalent for H3K4me3 and H3K27me3 exhibited high and low expression, respectively, with a lower level in LADs as compared to non-LADs (**Fig. 6c**). During neuronal differentiation, bivalent genes that resolved to the H3K4me3-monovalent state in LAD-loss regions exhibited a significant increase in their expression levels (**Fig. 6d, e)**. Interestingly, genes that remained bivalent in LAD-loss regions also increased their expression during neuronal differentiation (**Fig. 6f, g**), consistent with lamina-association playing a dominant role in their regulation. Importantly, genes that resolved to H3K4me3-monovalent state or remained bivalent in LAD-same and LAD-gain regions were still expressed at significantly lower levels (**Fig. 6e, g, Extended Data Fig. 9b, c**). Taken together, these data indicate that LAD changes occur early in the process of neuronal differentiation and support the notion that lamina-association contributes prominently to the lowly-expressed, poised state of bivalency.

### The lowly expressed state of lamina-associated bivalent genes is independent of H3K27me3

The removal of H3K27me3 resolves the bivalent chromatin to H3K4me3-monovalent state, which generally corresponds to increased gene expression. However, the observed increase in gene expression with H3K27me3 removal is often highly variable and many bivalent genes are not induced upon resolution of bivalency to H3K4me3 monovalency ^25,27^. We hypothesized that lamina association contributes to a transcriptionally repressive state independent of local levels of H3K27me3. To test this concept, we treated NPCs with Tazemetostat (EPZ-6438), a potent and selective inhibitor of EZH2 – the enzyme that catalyzes H3K27me3 – to globally reduce H3K27me3 levels (**Fig. 7a)**. As compared to DMSO vehicle control, treatment of NPCs for one week with Tazemetostat (0.5 µM, 1 µM) resulted in a near total loss of H3K27me3 peaks even at the lowest concentration of inhibitor (**Fig. 7b**).

**Figure 7.**
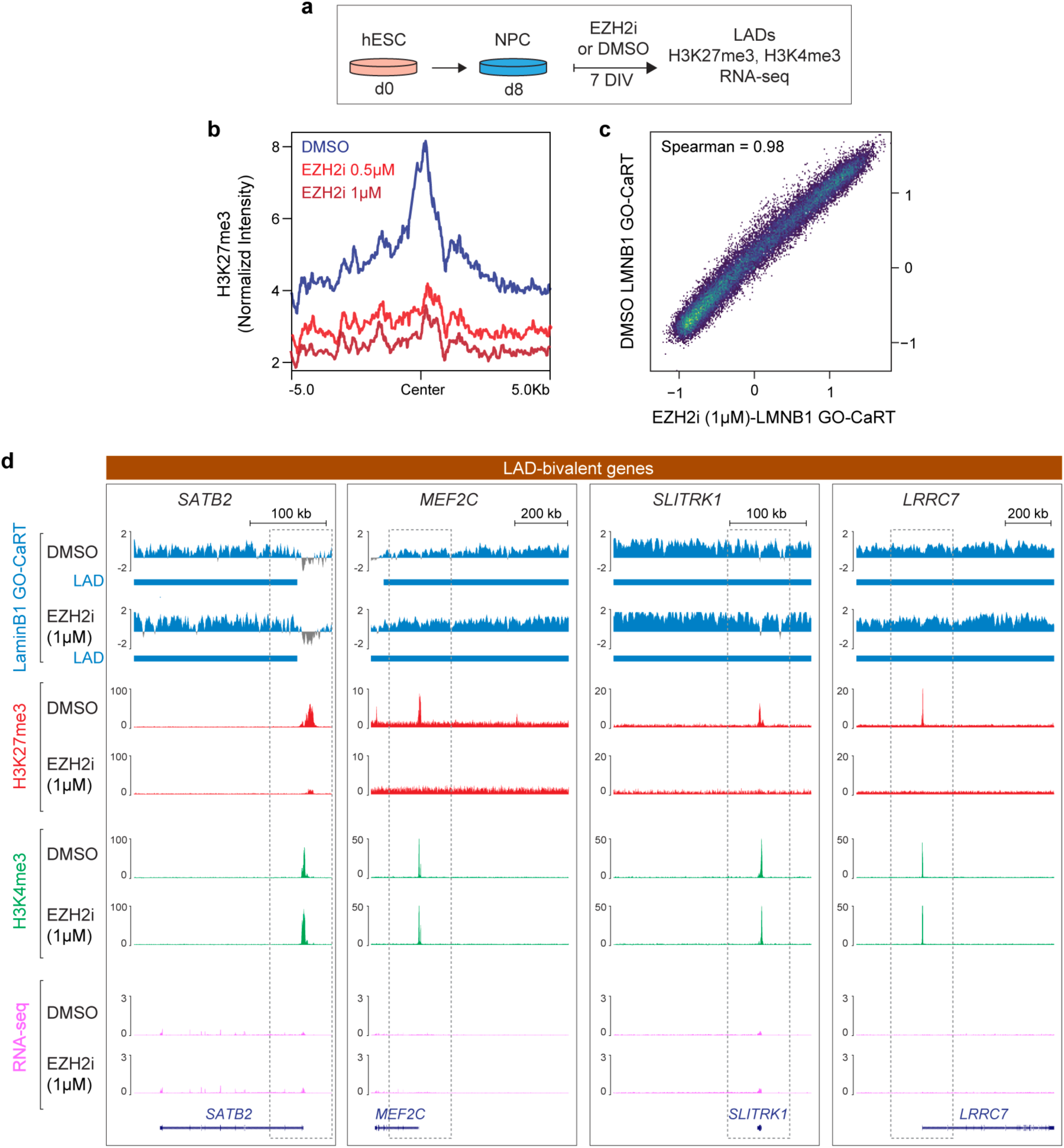
The low expression of lamina-associated bivalent genes is independent of H3K27me3. **a,** Schematic of EZH2 inhibition experiments in human ESC-derived NPCs. NPCs were treated with DMSO or EZH2 inhibitor, Tazemetostat/EPZ-6438 (referred to as EZH2i in figure panels) for 7 days in vitro (DIV) after which cells were harvested for LaminB1 GO-CaRT, CUT&RUN and RNA-seq. **b,** Normalized H3K27me3 levels in NPCs treated with DMSO, 0.5 μM and 1 μM of Tazemetostat (EZH2i) for 7 DIV. **c,** Genome-wide scatter plot showing Spearman correlation of LaminB1 GO-CaRT between DMSO and EZH2i (1 μM of Tazemetostat) treated NPCs. **d,** LaminB1 GO-CaRT, H3K27me3, H3K4me3 (CUT&RUN) and RNA-seq tracks at LAD bivalent genes, *SATB2*, *MEF2C*, *SLITRK1* and *LRRC7* in DMSO and EZH2i (1 μM Tazemetostat) treated NPCs. Dashed boxes show selective loss of H3K27me3 peaks and change of bivalent to H3K4me3-monovalent state at these loci with no apparent changes in transcription levels. Related data in Extended Data Fig. 10.

The LaminB1 GO-CaRT profiles of DMSO and Tazemetostat-treated NPCs were highly similar, indicating that EZH2 inhibition does not grossly disrupt LAD architecture of NPCs (**Fig. 7c, Extended Data Fig. 10a**). At bivalent genes, LaminB1 enrichment was also similar between DMSO and Tazemetostat-treated NPCs (**Extended Data Fig. 10b**). Although EZH2 inhibition led to derepression of all formerly bivalent genes, the expression level of those in LADs remained significantly lower (fold change= 2.37) than those in non-LAD regions (**Extended Data Fig. 10c**). In particular, *SATB2, MEF2C, SLITRK1,* and *LRRC7* remained in LADs and lowly expressed even after EZH2 inhibition (**Fig. 7d**). These results indicate that localization of bivalent genes at the lamina is not dependent on H3K27me3 and that their lowly expressed state is strongly reinforced by the repressive environment of the lamina.

In summary, we show that the dynamics of lamina and speckle–associated genome architecture regulates the activity of bivalent chromatin state and neurogenic transcriptional program during human cortical neurogenesis. Nuclear positioning near the lamina helps maintain the lowly expressed, poised transcriptional state of bivalent genes in stem/progenitor cells. During neurogenesis, extensive remodeling of spatial 3D genome relocates hundreds of neuronal genes from the lamina to speckles enhancing resolution of bivalent chromatin to the H3K4me3-monovalent state and strongly upregulates transcription (**Fig. 8 top**). Studies of neurogenesis in cell culture demonstrate that bivalent genes are embedded in the lamina and that lamina association maintains the lowly expressed state of bivalent genes independent of H3K27me3 (**Fig. 8 bottom**). This work builds a scientific paradigm in which spatial genomic location in relation to subnuclear compartments critically informs the activity of local chromatin state and gene regulation.

**Figure 8.**
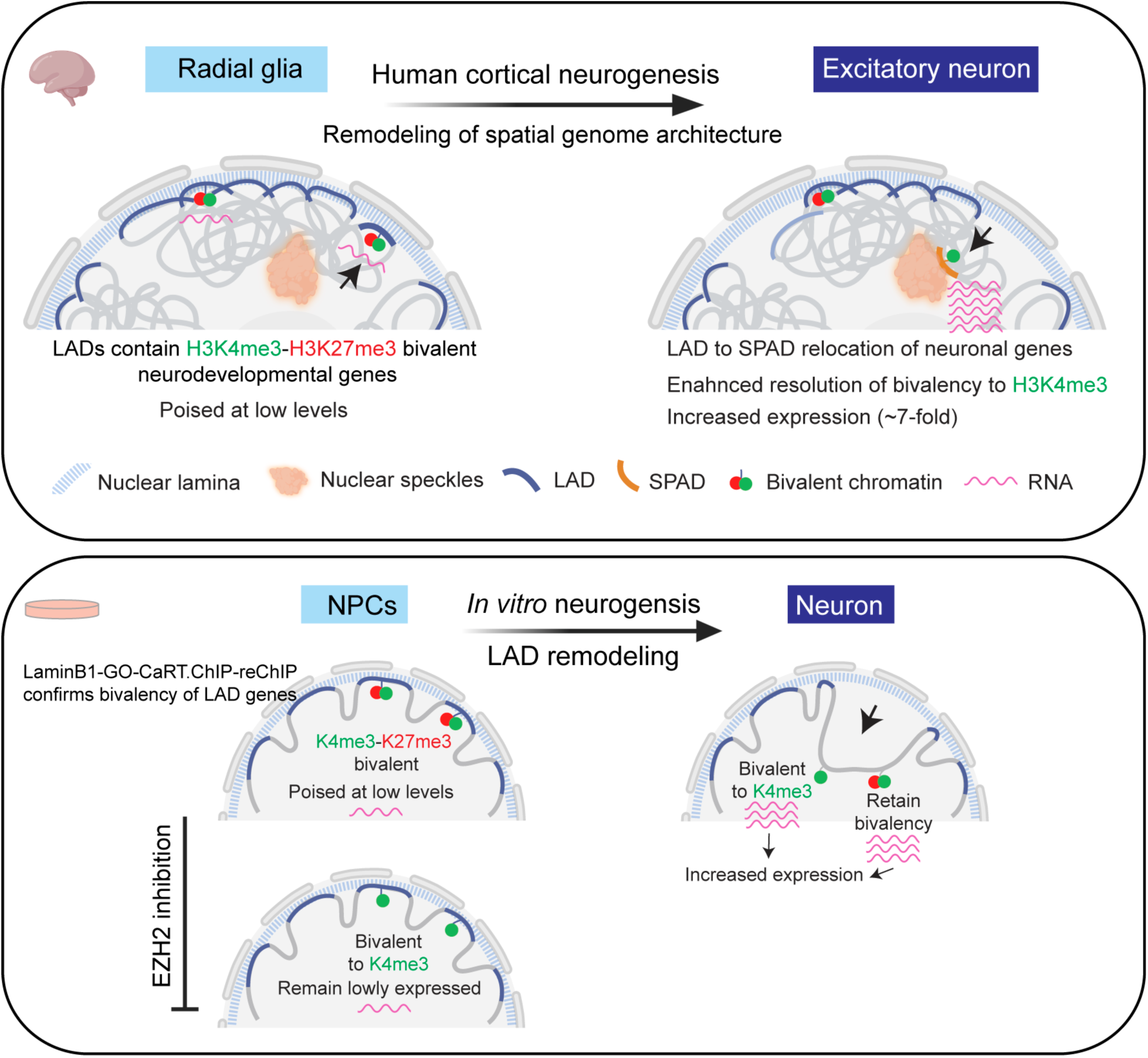
A summary of how spatial 3D genome dynamics controls the activity and transcriptional output of bivalent chromatin during human neurogenesis. **(top)** In neural progenitor cells (Radial glia) of the developing human brain, neurodevelopmental genes marked by H3K4me3-H3K27me3 bivalent chromatin are enriched at the lamina and are expressed at very low levels. During cortical neurogenesis (RG to EN), extensice remodeling of the 3D genome relocates hundreds of neuronal genes from the lamina to speckles. Lamina-to-speckle translocation enhances resolution of bivalent chromatin to the H3K4me3-monovalent state and strongly increases transcription. **(bottom)** Cell culture models of human neurogenesis confirm the bivalent state of genes embedded in the lamina and provide further evidence that lamina association – and not the presence of H3K27me3– is the dominant factor in maintaining the lowly expressed, poised transcriptional state of bivalent genes.

## Discussion

Our knowledge of the organization and dynamics of spatial 3D genome organization around subnuclear compartments and how it regulates developmental gene expression remains scarce. By developing methods that enable the study of specific cell types isolated directly from rare samples of midgestational human cortex, we were able to generate high-resolution maps of LADs and SPADs, discovering critical relevance of their dynamics during the birth of neurons in the neocortex. We used these maps to investigate the role of lamina and speckle– associated genome organization and how it interacts with bivalent chromatin state to regulate lineage-specific gene expression. This analysis yielded broader insights into how genome positioning with respect to subnuclear compartments impacts transcriptional output of the bivalent chromatin state – a key feature of developmentally important genes. Additional studies in iPSC/ESC-derived NPCs provided experimental evidence that proximity to the nuclear periphery – and not the presence of H3K27me3 – is the dominant factor in maintaining the lowly expressed, poised state of bivalent genes embedded in the lamina. Together, our work reveals a critical role of spatial genomic localization in relation to subnuclear compartments in determining lineage-specific gene expression during brain development, highlighting the importance of 3D spatial genome organization in epigenomic transcriptional regulation.

### Large-scale reorganization of lamina-associated genome during human cortical neurogenesis

We found that during human cortical neurogenesis, the remodeling of LAD architecture is extensive, involving ∼19% of the whole genome and encompassing Mb-scale genomic regions. Such large-scale changes to LAD architecture have not been observed in cell culture models of differentiation. For example, 73-87% of LAD coverage is shared between mouse embryonic stem cells (mESCs) and ESC-derived NPCs and astrocytes ^19^. Similarly, the majority of the LADs (94%) remain unchanged during *in vitro* differentiation of proliferative myoblasts to myotubes ^33^. *In vitro* differentiation of mESCs into cardiomyocytes also does not lead to major changes in LAD architecture ^34^. Although our *in vitro* models of human neurogenesis recapitulated key aspects of *in vivo* LAD dynamics, the remodeling of LAD architecture was less extensive. In addition to differences related to the potential effects of cell culture, one possibility is that the *in vivo* developmental context is required for large-scale LAD remodeling. Our result corroborates with previous study of LADs in mouse preimplantation embryo development during which major changes to LAD organization are observed ^35^. It is also possible that longer time intervals of development along the lineage being studied are required for large-scale LAD remodeling. These results underscore the importance of studying genome architecture dynamics in cells that develop *in vivo*.

### Lamina-to-speckle 3D genome dynamics regulates neuronal gene expression

Of the 1240 genes that detached from the lamina during cortical neurogenesis, nearly 56% (696 genes) became associated with nuclear speckles. This degree of apparent movement of hundreds of neuronal genes from LADs to SPADs indicates that genomic positioning relative to nuclear speckles is highly dynamic during development. While the molecular mechanisms that underlie this movement of genes from LADs to SPADs remains to be determined, developmentally important transcription factors could be involved in targeting genes to speckles. Recent work provided evidence that transcription factors can target genes to speckles for enhanced transcription ^36^. Furthermore, the finding that genes near the speckles exhibit both increased gene expression and greater isoform diversity suggests that genome positioning near speckles may facilitate efficient coupling of transcription and splicing. Indeed, recent work has shown that genes located near the speckles exhibit increased co-transcriptional splicing and recruitment of a pre-mRNA to speckles is sufficient to drive increased mRNA splicing levels ^37^. This may be particularly important for neuronal genes which are generally long and exhibit higher isoform diversity ^38^. Thus, LAD-to-SPAD switching is a key component of lineage-specification and neuronal differentiation during human brain development.

Nuclear speckles have long been proposed to act as physical hubs for active genes ^4,13,15^. Recent in-depth genomic analysis has revealed their important function in 3D genome organization and transcriptional regulation ^10,16,17^. TSA-seq revealed a largely similar genome organization around speckles across four human cell lines (H1-ESCs, HFF, HCT-116 and K562)^20^. While this observation also holds true for SPADs in proliferative neural progenitors (RG, IPC) in the human brain, we find that the SPAD architecture of postmitotic neurons (EN) is particularly distinct. This indicates that the SPAD architecture of cells *in vivo* is dynamic and exhibits key differences between cells that are proliferating (RG, IPC) and those that have exited the cell cycle (EN). We observed that SPADs in EN are enriched in genes critical to neuronal function. Many neurological disorders exhibit defects in splicing of neuronal genes ^39,40^ and we previously found that SPADs in the human cortex are enriched in loci associated with genetic risk for schizophrenia^10^. Thus, nuclear speckles may be of particular importance to neuronal 3D genome organization and function.

### 3D spatial genome positioning is a key determinant of transcriptional regulation associated with the bivalent chromatin state

The H3K27me3-H3K4me3 bivalent chromatin state is a key feature of developmentally important genes that was first described in ESCs ^23,24^. While many bivalent genes are highly activated upon differentiation-associated resolution to the H3K4me3-monovalent state, many other resolved genes are not induced during differentiation ^25,27^, suggesting that additional factors contribute to bivalency-associated transcriptional regulation. Our work moves beyond the linear interpretation of chromatin state and illustrates how spatial 3D genome organization can exert control over the activity and transcriptional output of bivalent chromatin. We find that bivalent genes that reside in LADs – both in RG and iPSC-derived NPCs – are expressed at much lower levels as compared to the bivalent genes in the rest of the genome, suggesting that the lamina contributes an additional layer of transcriptional repression. This may be particularly important to prevent any unscheduled activation of developmental genes, thereby conferring robustness to the system. Consistent with this notion, during neurogenesis, bivalent genes that resolve to the H3K4me3-monovalent state but remain in LADs are only moderately induced, while those that detach from the lamina exhibited ∼2-4-fold increase in transcription. Moreover, genes that retain bivalency but become dissociated from the lamina during the differentiation of NPCs to neurons are also transcriptionally upregulated. These findings support the idea that localization near the lamina confers a lowly expressed ‘poised’ transcriptional state to genes marked by bivalent chromatin in stem/progenitor cells, and importantly provide a more nuanced interpretation of the functional impacts of bivalency resolution.

The lowly expressed, poised state of bivalent genes in LADs could be attributed to the transcriptionally repressive nature of H3K27me3 modification, which is enriched at LAD borders ^10,11,41^. Our data do not suggest such a role for H3K27me3 in the low expression of LAD-resident bivalent genes. Despite global loss of H3K27me3 via pharmacological inhibition of EZH2, genome-lamina association is not grossly perturbed and the genes in LADs remain lowly expressed. These findings indicate that the lamina provides other activities that help maintain a low but ‘poised’ state of expression for bivalent genes.

Our data show that the resolution of bivalency to the H3K4me3-monovalent state is variable across the genome and is influenced by spatial genomic location with respect to subnuclear compartments. During RG to EN differentiation, ∼30-35% of bivalent genes resolve to the H3K4me3-monovalent state when remaining at the lamina. When bivalent genes dissociate from the lamina and relocate to speckles during RG to EN differentiation, ∼85% resolve to the H3K4me3-monovalent state and exhibit ∼7-fold increase in transcription. This suggests that localization to the lamina may serves as a barrier to resolution of bivalency to the H3K4me3 active state by restricting access of transcription factors and chromatin remodelers linked to the resolution of bivalency ^7,42^. On the other hand, the enhanced resolution of bivalent to H3K4me3-monovalent state associated with lamina-to-speckle relocation may be a consequence of transcriptional induction, but it is also plausible that the speckles contain factors that help resolve bivalency. Thus, 3D spatial location relative to specific subnuclear compartments more accurately informs the transcriptional consequences of the local chromatin state.

### Exploring 3D spatial genome organization using improved ‘cell-type specific’ GO-CaRT

We previously used GO-CaRT to study LADs and SPADs in prenatal brain tissues from various mammals including humans, and these bulk specimen studies provided important insights about conservation of genome architecture across species and differences between distinct brain regions ^10^. Studying specific cell lineages isolated by antibody-based sorting poses downstream problem of unintended pA/G-MNase targeting while performing GO-CaRT. This methodological challenge was addressed in this study. We combined Fab blocking with GO-CaRT to enable studies of LAD and SPAD dynamics within specific cell types of a commonly studied neurogenic lineage in prenatal human brain specimens. This cell type-specific atlas of LADs and SPADs thus provides the first analysis of this aspect of 3D genome architecture within a developmental lineage as it occurs *in vivo.* Fab blocking avoids the concerns of unintended targeting of pA/G-fused MNase (or similar pA/G-fused enzymes e.g., pA/G-Tn5) to antibodies used in FACS/FANS, particularly those against nuclear antigens. With the development of this ‘cell-type specific’ GO-CaRT method, we are better positioned to explore the spatial 3D genome organization relative to the lamina, speckles and other less-explored subnuclear compartments (e.g., nucleolus, PML bodies, Cajal bodies etc.) in various cellular contexts *in vivo*. Moreover, the GO-CaRT-sequential ChIP approach developed here should facilitate investigations that further advance our understanding of the interplay between spatial genome organization and local chromatin state.

In conclusion, our work demonstrates the large-scale, highly dynamic nature of spatial 3D genome architecture in neurodevelopment, far beyond what has been observed in cell culture studies. Furthermore, the use of these maps provides critical information for the interpretation of local chromatin state, leading to a model wherein spatial location inside the nucleus dominates the regulation of bivalent chromatin loci. More broadly, these data establish a new paradigm for understanding epigenomic transcriptional regulation, where genome’s spatial organization is combined with local chromatin state to gain better insights into the mechanisms of gene expression control.

## Methods

### Tissue collection and processing

Human brain tissue samples (GW17 and GW20) were collected from de-identified donors with previous patient consent in strict observance of the legal and institutional ethical regulations. All protocols were approved by the Human Gamete, Embryo and Stem Cell Research Committee (GESCRC) and Institutional Review Board (IRB) at the University of California, San Francisco (UCSF). Primary human brain tissue was collected and processed as previously described ^43^. Briefly, cortical tissue was collected in artificial cerebrospinal fluid (ACSF) containing 125 mM NaCl, 2.5 mM KCl, 1 mM MgCl_2_, 1 mM CaCl_2_ and 1.25 mM NaH_2_PO_4_ under a stereotaxic dissection microscope (Leica). Tissue samples were cut into small pieces and snap-frozen by placing the tissue on a strip of aluminum foil in pre-chilled 2-methylbutane on dry ice for ∼10 min. Samples were stored in cryovials at −80 °C. For individual germinal zone (GZ) and cortical plate (CP) cultures, samples were dissected in artificial CSF to separate GZ from the CP before dissociation. Tissue pieces were then placed in a pre-warmed (at 37 °C) solution of papain (Worthington) and incubated at 37 °C. After approximately 60 min of incubation, tissue was triturated following the manufacturer’s protocol, and samples were spun through an ovomucoid gradient to remove debris. The dissociation media was removed, and cells were resuspended in NES media ((DMEM/F12 (gibco), 1:1,000, B27 (Invitrogen), 1:100 N2 (Life Technologies), 20 ng/ml of FGF (PeproTech), 20 ng/ml of EGF (PeproTech), 20 µg/ml of insulin (Thermofisher), 5 ng/ml of BDNF (PeproTech) and 10 µM ROCK inhibitor Y-27632 (Selleckchem)). Cells were plated in Matrigel (Corning) coated (1µg/mL) 8-well chambered coverglass (Thermofisher) and allowed to attach for 24 hours at 37°C.

### Cell culture maintenance

Human pluripotent stem cells (WTC11-iPSC and H1-ESC) were cultured in Matrigel-coated 6-well plates in StemFlex media (Gibco). Media was supplemented with 10 µM ROCK inhibitor Y-27632 (ROCKi) for the first 16-20 hours of culture and was subsequently removed. Cells were fed fresh media every day until they reached 80-90% confluency at which point they were passaged as clumps using ReLeSR (Stemcell Technologies). HEK293T cells were cultured in DMEM (Gibco) supplemented with 10% FBS and 1X antibiotic/antimycotic mixture (Gibco).

### Generation of NPCs from human pluripotent stem cells

Human iPSC line WTC11 (Kreitzer et al. 2013) ^44^ was differentiated into NPCs as described in Ciceri et al. 2024 ^32^ with modifications. Cells were dissociated with Accutase (Thermofisher) and seeded on Matrigel-coated plates at a density of 150,000 cells/cm^2^ in StemFlex supplemented with 10 µM ROCKi. The following day (day 0), cells were washed in PBS and fed with neural induction medium (NIM: DMEM/F12 Glutamax (Gibco) containing 1% ITS-G (Gibco) and 200 µM L-ascorbic acid 2-phosphate (Santa Cruz). On days 0-2, cells were fed daily with NIM supplemented with 100 nM LDN-193189 (SelleckChem), 10 µM SB431542 (SelleckChem), and 2 µM XAV939 (Cayman). On days 3-9, NIM was supplemented with LDN-193189 and SB431542 only. From day 10-19, cells were fed daily with NPC maturation medium (1:1 mixture of DMEM/F12 Glutamax and Neurobasal-A supplemented with 1% N-2 and 2% B-27 minus vitamin A (Gibco). On day 20, NPCs were either harvested for analysis or further differentiated into neurons.

Human H1-ESC line was differentiated into NPCs using the StemXVivo Neural differentiation kit (R&D systems) following manufacturer’s instructions. Briefly, ESCs were dissociated using ReLeSR and plated as small clumps in a Matrigel-coated 6-well plate in StemFlex media with 10 µM ROCKi. Next day, the media was removed and replaced with NPC differentiation Media (Day 0). Cells were fed fresh NPC differentiation media for seven days. On Day 8, NPCs were dissociated and continued to passage as NPCs or used for experimental analyses.

### Neuronal differentiation of iPSC-derived NPCs

On day 20, NPCs were dissociated with Accutase and seeded at a density of 275,000 cells/cm^2^ on plates coated with 100 µg/mL poly-L-ornithine, 100 µg/mL poly-D-lysine, 10 µg/mL laminin and 10 µg/mL fibronectin (Sigma) in neuronal differentiation medium (Neurobasal-A supplemented with 2% B-27, 2 mM L-glutamine (Gibco), 200 µM L-ascorbic acid 2-phosphate, 10 µM dibutyryl-cyclic AMP (Sigma), 10 ng/mL BDNF (Peprotech) and 10 ng/mL GDNF (Peprotech). Neuronal differentiation media was supplemented with 10 µM DAPT (MedChemExpress) and cells were fed by 50% medium exchange every 2 days. After one week of neuronal differentiation, cells were harvested for downstream experimental analyses.

### EZH2 inhibition in NPCs

On day 8, H1-ESC-derived NPCs were passaged using Accutase and plated into a Matrigel-coated 6-well plate in NPC differentiation media supplemented with 10 µM ROCKi. Next day, the media was removed and fresh media containing EZH2 inhibitor Tazemetostat (EPZ-6438, Selleckchem, 0.5 μM, 1 μM) or DMSO was added to the cells. Cells were fed fresh media containing the inhibitor (or DMSO) every day for one week at which point they were harvested for downstream experimental analyses.

### Nuclei Isolation and FANS

Snap-frozen cortical tissue was cut into small pieces and transferred to a pre-chilled 7-ml Wheaton™ Dounce Tissue Grinder containing 5 ml nuclei extraction buffer (NEB: 10 mM HEPES pH 7.4, 25 mM KCl, 5 mM MgCl2, 0.25 M sucrose, 0.1% Triton X-100, 1x Halt protease inhibitor cocktail (Thermofisher)). RiboLock (Thermofisher) was added to all the buffers to preserve RNA integrity. While still on ice, the tissue was dissociated with 5-6 strokes of loose pestle (A) followed by 8-10 strokes of tight pestle (B), until no tissue pieces were visible. The sample was incubated on ice for ∼5 min after which it was transferred to a pre-chilled 15-ml conical tube. The sample was centrifuged at 500g for 10 min at 4°C. The supernatant was removed, and the nuclear pellet was resuspended in 10-ml NEB without Triton X-100. The homogenate was passed through a 40μ-strainer into a 50-mL conical tube. Formaldehyde (Thermofisher) was added to a final concentration of 0.1% and the sample was incubated for 2 min at room temperature with rotation. Glycine was added to a final concentration of 75 mM to quench the reaction. BSA was added to a final concentration of 1% and the sample was centrifuged at 500g for 10 min at 4°C. The supernatant was discarded, and the nuclear pellet was resuspended in 1 mL staining buffer (PBS + 1% BSA). The nuclei were filtered through a 40-µ strainer into a 1.5-mL low-bind microcentrifuge tube and counted under the microscope. The nuclei were pelleted at 500g for 10 min at 4°C and resuspended in 100-150 μl of staining buffer. Antibody staining was carried out for 1 hour at 4°C on rotation with the following antibodies: PAX6-Alexa Fluor 488 (BD Biosciences,1:20 dilution), EOMES-PE-Cy7 (Invitrogen, 1:20 dilution), and SATB2-Alexa Fluor 647 (Abcam, 1:100 dilution). After staining, 900 μl of staining buffer was added and the samples were centrifuged at 500g for 10 min at 4°C. The nuclear pellet was resuspended in 1 mL of staining buffer and centrifuged at 500g for 10 min at 4°C. Nuclei were resuspended in 1-2 mL of staining buffer (depending upon the starting material and yield) and filtered into a FACS (BD) tube. DAPI was added at 1μg/ml just before FANS. AbC™ Total Compensation capture beads (Thermofisher) were used for generating single color compensation controls. FANS was conducted on BD FACS Aria II Cytometer using a 70-μm nozzle. Sorted nuclei were collected in 5-ml tubes containing 300-500 μl of collection buffer (PBS + 5% BSA) and RNasin Plus RNase inhibitor (Promega). Sorted nuclei were collected by centrifuging at 500g for 10 min at 4°C and processed for downstream analyses (RNA-seq, Fab blocking and GO-CaRT/CUT&RUN).

### Development of Fab blocking for antibody-based chromatin profiling

HEK293T cells were harvested by trypsinization. Cells were fixed in 0.1% Formaldehyde for 2 minutes at room temperature, followed by quenching in 0.1M Glycine (prepared in wash buffer). Cells were resuspended in 1 ml wash buffer (20 mM HEPES-KOH, pH 7.5, 150 mM NaCl, 0.1% BSA, 0.5 mM spermidine (sigma) and 1× Halt protease inhibitor cocktail)) and centrifuged at 300g for 5 min. This step was repeated for a total of two washes. Cells were resuspended in wash buffer and aliquoted at 100 μl/tube into 0.5 ml PCR tubes (∼100k cells per experimental condition). Next, 8 μl of BioMagPlus concanavalin A beads (Polysciences) activated in binding buffer (20 mM HEPES-KOH, pH 7.9, 10 mM KCl, 1 mM CaCl2 and 1 mM MnCl2) were added to each sample and rotated on a nutator for 10 min at room temperature. The cells were placed on a magnet to clear and the liquid was removed. Cells bound to beads were resuspended in 50 μl cell permeabilization buffer (20 mM HEPES, pH7.5, 0.1 mM CaCl2, 3 mM MgCl2, 100 mM KCl, and 0.05% Digitonin (MilliporeSigma) and 1X Halt protease inhibitor cocktail added fresh)) and incubated for 30 min at room temperature on a nutator. The supernatant was removed, and the beads were resuspended in 50 μl of antibody binding buffer (wash buffer + 2 mM EDTA + 0.025% digitonin) containing an antibody against H3K27me3 (Cell Signaling Technologies, 1:100 dilution of 1 mg/ml). IgG (Cell Signaling Technologies, 1:100 dilution) was included as a negative control. The samples were incubated overnight at 4°C on a nutator. Next day, the supernatant was removed, and the beads were washed two times with 200 μl wash buffer containing 0.025% digitonin (Wash-Dig). Beads were resuspended in 50 μl of Wash-Dig containing monovalent anti-rabbit Fab fragments (Jackson Immuno) and incubated for 30 min at 4°C. We tested the following Fab dilutions: 1:20, 1:50, 1:100, 1:250, 1:500, 1:1000, 1:10,000, 1:50,000 and 1:100,000. We also tested three incubation times: 5 min, 15 and 30 min. Dilutions 1:250 and lower and incubation times 5 min and below did not result in efficient blocking by the Fab fragments. Following two washes with WaB-Dig, beads were resuspended in 50 μl of WaB-Dig containing pA/G-MNase (purified from addgene plasmid at Macro Lab UC Berkley) and incubated at 4°C on nutator for 1 hour. The supernatant was removed, and the beads were washed two times with WaB-Dig. Beads were resuspended in 100 μl of WaB-Dig containing 2 mM CaCl2 to activate pA/G-MNase. The digestion was carried out for 30 minutes in chilled metal blocks on ice. Digestion was stopped by adding 100 μl of 2XSTOP (200 mM NaCl, 20 mM EDTA, 4 mM EGTA, 50 μg/ml of RNAseA, 40 μg/ml of glycogen and 10 pg/ml of heterologous DNA). Samples were incubated at 37°C for 30 min to release pA/G-MNase cleaved fragments. The contents of the tube were transferred to 1.5-ml eppendorf tube containing 2 μl of each 10% SDS and Proteinase K (20 mg/mL). The samples were incubated at 55 °C for 1 hour to reverse crosslinking. DNA was extracted using Phenol: Chloroform extraction method. Purified DNA fragments were analyzed by Tapestation High Sensitivity D1000 (Agilent).

To demonstrate that Fab blocking allows subsequent binding of another antibody for GO-CaRT/CUT&RUN analyses, we incubated the Fab blocked H3K27me3 samples with an antibody against LaminB1 protein. Briefly, following Fab blocking of H3K27me3, beads were resuspended in 50 μl of WaB-Dig containing LaminB1 antibody (Abcam, 1:100 of 1 mg/mL). The sample was incubated at 4 °C on nutator for 2 hours. Following two washes with WaB-Dig, the steps of pA/G-MNase binding, digestion and DNA extraction were carried out as described above. Purified DNA fragments were analyzed by Tapestation High Sensitivity D1000 (Agilent).

### Fab blocking followed by GO-CaRT and CUT&RUN analyses

Nuclei sorted by FANS were resuspended in PBS containing 1% BSA and divided into 100 μl aliquots in 0.5-ml PCR tubes. Next, 8 μl of activated BioMagPlus concanavalin A beads were added to each sample and rotated for 10 min at room temperature. The samples were placed on a magnetic stand to remove the liquid. To block antibodies used in FANS, nuclei-bound to concanavalin A beads were resuspended in 50 μl wash buffer (20 mM HEPES-KOH, pH 7.5, 150 mM NaCl, 0.1% BSA, 0.5 mM spermidine and 1× Halt protease inhibitor cocktail) containing anti-rabbit and anti-mouse (Jackson Immuno) monovalent Fab fragments at 1:30 dilution. The samples were incubated on a nutator at 4 °C for 30 min. The samples were placed on a magnetic stand to remove the liquid. Beads were resuspended in 200 μl wash buffer to remove excess Fab fragments. The beads were resuspended in 50 μl antibody binding buffer (wash buffer + 2 mM EDTA) containing primary antibody of interest for GO-CaRT/CUT&RUN. The following antibodies were used: anti LaminB1 antibody (Abcam, 1:100 dilution) for LaminB1 GO-CaRT, anti-SON antibody (Atlas Antibodies, 1:100 dilution) for SON GO-CaRT, anti-H3K27me3 (Cell Signaling Technologies, 1:50 dilution) and anti-H3K4me3 (Cell Signaling Technologies, 1:100 dilution) for CUT&RUN. An IgG control was run in parallel for all experiments (Cell Signaling Technologies, 1:100 dilution). Antibody incubation was carried out overnight on a nutator at 4 °C. The next day, samples were briefly spun and placed on a magnetic stand to remove the liquid. Samples were washed two times with 200 μl of wash buffer. Beads were resuspended in 50 μl wash buffer containing pA/G-MNase and rotated for 1 hour at 4 °C. Samples were briefly spun and placed on a magnetic stand to remove the liquid. Samples were washed two times with 200 μl of wash buffer. Beads were gently resuspended in a 100 μl of wash buffer containing 2 mM CaCl2 (to activate pA/G-MNase) and placed in a pre-chilled metal blocks sitting in ice. Digestion was carried out for 30 min and stopped by adding 100 μl of 2XSTOP (200 mM NaCl, 20 mM EDTA, 4 mM EGTA, 50 μg/ml of RNAseA, 40 μg/ml of glycogen and 10 pg/ml of heterologous DNA). Samples were incubated at 37 °C for 20 min to release pA/G-MNase cleaved fragments. 2 μl SDS (10%) and 2 μl Proteinase K (20 mg/ml) was added to each sample and incubated at 55 °C for 1 hour. DNA was extracted using the phenol–chloroform extraction method. Purified DNA fragments were analyzed by Tapestation High Sensitivity D1000 (Agilent).

### LaminB1 GO-CaRT.ChIP-reChIP

NPCs derived from iPSC line WTC11 were harvested using Accutase and the cell pellet was resuspended in 1 mL ice cold PBS (Gibco). The cells were centrifuged at 500g for 3 min at 4 °C and the cell pellet was resuspended in 1 ml of nuclei isolation buffer (NIB: 10 mM HEPES-KOH, pH 7.9, 10 mM KCl, 0.1% NP40, 0.5 mM spermidine and 1× Halt protease inhibitor cocktail) and incubated for 10 min on ice. Nuclear pellet was collected by centrifuging at 600g for 3 min at 4°C and again resuspended in 1 ml of NIB to wash. Nuclear pellet was collected by centrifugation at 600g for 3 min and resuspended in 100 μl of NIB.

LaminB1 GO-CaRT was performed as described above, and LaminB1-enriched chromatin from the supernatant was used for subsequent sequential (2-step) ChIP. LaminB1 GO-CaRT was performed in twelve 0.5 ml PCR tubes, each containing ∼500,000 nuclei (total ∼6 million nuclei) in 100 μl of NIB. For antibody and pA/G-MNase binding steps, the solutions were scaled up from 50 μl to 100 μl. The reaction was stopped using 2XSTOP containing LaminB1 blocking peptide (Abcam) at 15 μg/ml followed by incubation at 37 °C for 15 min. The samples from 12 PCR tubes were pooled together and centrifuged at 14,000g for 5 min at 4 °C. Supernatant containing LaminB1-associated DNA fragments was divided into 300 μl aliquots for 1^st^ ChIP. One aliquot was kept as input and stored at −20 °C for later use. For each ChIP, we used LaminB1 GO-CaRT supernatant from ∼750k nuclei. Antibodies for LaminB1 (1:100), H3K4me3 (1:100), H3K27me3 (1:50) and IgG control (1:100) were added and the samples were incubated at 4 °C overnight with rotation. The next day, Protein A dynabeads (Thermofisher) were equilibrated by washing them two times in 1 ml wash buffer containing 0.05% Tween-20 (Wash + Tween). Beads were resuspended in Wash + Tween in the original volume of beads. 15 μl of equilibrated beads were added to each ChIP reaction and the samples were incubated at 4°C for 1 hour. Samples were washed two times with 1 ml Wash + Tween by rotating for 5 min at room temperature. The tubes were placed on the magnet to remove the supernatant. At this point, controls for 1^st^ ChIP (LaminB1 GO-CaRT.LaminB1 ChIP and LaminB1-GO-CaRT.IgG ChIP) have finished processing and these samples were resuspended in 200 μl of Wash + Tween and stored at −20 °C for DNA extraction later. For H3K4me3 and H3K27me3 1^st^ ChIP samples, the chromatin was eluted by resuspending the beads in 150 μl of Wash + Tween containing the corresponding blocking peptides (10 μg/mL) for H3K4me3 (EpigenTek) and H3K27me3 (EpigenTek), respectively. The samples were incubated at 4 °C for 2 hours. For H3K4me3 and H3K27me3, the 1^st^ ChIP was carried out in two tubes, so the total volume of eluted chromatin was 300 μl. The samples were placed on the magnet and the supernatant containing eluted chromatin from 1^st^ ChIP was transferred to a new 1.5-ml tube. A 100 μl aliquot of eluted chromatin was saved as 1^st^ ChIP input. To the remaining 200 μl of chromatin, an antibody of interest for the 2^nd^ ChIP (H3K27me3 if 1^st^ ChIP was done with H3K4me3 or H3K4me3 if 1^st^ ChIP was done with H3K27me3) was added and the samples were rotated at 4 °C overnight. Next day, 15 μl of equilibrated Protein A dynabeads were added to each 2^nd^ ChIP sample and incubated at 4 °C for 1 hour. Samples were washed two times with 1 ml Wash + Tween by rotating for 5 min at room temperature. The tubes were placed on the magnet to remove the supernatant and the beads were resuspended in 200 μl of Wash + Tween. To each sample (including samples from 1^st^ ChIP and inputs), 2 μl of SDS (0.1% final) and 2 μl of 20 mg/ml Proteinase K were added. The samples were vortexed and incubated at 55 °C for 1 hour. DNA was extracted using the phenol–chloroform extraction method.

### Library preparation and sequencing

Sequencing libraries for GO-CaRT, CUT&RUN and LaminB1-GO-CaRT.ChIP-reChIP were generated using KAPA HyperPrep Kit (KAPA, KK8504) following the manufacturer’s instructions. The libraries were amplified for 12–14 PCR cycles. DNA fragments were quantified by Tapestation D1000 (Agilent) and Qubit high sensitivity dsDNA (Invitrogen) assays, pooled, and sequenced by 150-bp paired-end sequencing on NovaSeq 6000 or NovaSeq X Plus systems (Illumina).

### CUT&Tag

The CUT&Tag was implemented following the protocol described in Kaya-Okur et al. ^45^ and Zhang et al. for profiling LaminB1, H3K4me3 and H3K27me3 during neuronal differentiation from iPSC-derived NPCs. Briefly, iPSC-derived NPCs and neurons were harvested by scraping in 1x Nuclei Isolation Buffer (NIB), triturated, and incubated on ice for 10 minutes. Cells were then centrifuged at 500g for 5 min at 4°C. Nuclei were resuspended in PBS before fixation by 0.1% formaldehyde for 2 min. Fixation was quenched by addition of 75 mM glycine for 2 min incubation on ice. Nuclei were centrifuged and resuspended in wash buffer 1 (20 mM HEPES pH 7.4, 150 mM NaCl, 0.5 mM spermidine). For each reaction, 100,000 cells were counted for binding to activated Concanavalin A-coated magnetic beads and incubated with primary antibodies overnight on a nutator at 4°C. On day 2, samples were placed on a magnetic stand and washed by wash buffer 1 for three times. Beads were resuspended in Wash Buffer 1 containing secondary antibody at 1:100 dilution and nutated for 1 hour at 4°C. After clearing on magnetic stand, beads were washed twice with Wash Buffer 1, resuspended in wash buffer 2 (20 mM HEPES pH 7.4, 300 mM NaCl, 0.5 mM spermidine) containing protein A-fused preloaded Tn5 transposase at 1:100 dilution, and nutated for 1 hour at 4°C. Samples were then washed twice with wash buffer 2, resuspended in 50 µl wash buffer 1 containing 10 mM MgCl_2_ for subsequent transposition at 37°C for 1 hour. Tagmentation reaction was quenched by addition of 1.7 μl 0.5M EDTA, 0.5 μl 10% SDS, and 0.5 μl 20 mg/mL proteinase K for 1 hour at 55°C. DNA was phenol-chloroform extracted and resuspended in 1x TE buffer (pH 8.0). DNA was PCR amplified by adding 2 µl each of 10 µM universal i5 primer and uniquely barcoded i7 primer ^46^, along with 25 µl NEBNext High-Fidelity 2X PCR Master Mix (NEB). PCR products were size-selected with SPRIselect beads (Beckman Coulter) to recover DNA fragments over 75bp according to the manufacturer’s instructions. Eluted DNA fragments were quantified by Tapestation D1000 (Agilent) and Qubit high sensitivity dsDNA (Invitrogen) assays, pooled, and sequenced by 150-bp paired-end sequencing on a NovaSeq 6000 system (Illumina).

### GO-CaRT data processing and domain calling

Sequencing reads were first trimmed to remove adapters and low-quality sequences using Trim Galore (https://www.bioinformatics.babraham.ac.uk/projects/trim_galore/) with default parameters. Then trimmed reads were mapped to the human reference genome (hg38) using Bowtie2 (v2.3.4.4). PCR duplicates were removed using Picard tools (http://broadinstitute.github.io/picard/). Properly paired reads with high mapping quality (MAPQ score > 30) were kept for further analysis. For visualization on the UCSC Genome Browser, the final bam files of LaminB1 / SON were converted to bigwig files using bamCompare of deepTools (v3.4.1) with default parameters with IgG as the control bam files. For the visualization of histone modifications, bigwig files were generated using bamCoverage of deepTools (v3.4.1) with default parameters. For domain calling, the genome was binarized into 10kb bins and LADs and SPADs were called using SEACR (v1.3) that is specifically designed to call broad enriched regions as per instruction. Each target data bedgraph was normalized by the corresponding IgG control, where the peak of the curve was set as the threshold during the domain calling. LADs/SPADs overlapped with blacklist ^47^ were discarded.

### Chromosome-specific LAD/SPAD calling

For each biological replicate, all aligned reads passing the QC standards on each chromosome were included in the analysis. LADs/SPADs identified in two replicates were plotted in a by-chromosome manner using R package ggplot2.

### Annotations

Promoters were defined as ±1 kb around the TSS of the longest transcript of each gene (Ensemble release ^48^). Bivalent promoters were those that contained peaks for both H3K4me3 and H3K27me3.

### Analyses of LAD reorganization

LAD-same regions were identified based on their consistent presence in all stages of neurogenic lineage (RG, IPC and EN). LAD-loss regions were identified based on their presence in RG and IPC and loss in EN. LAD-gain regions were identified based on their presence in EN and absence in RG and IPC. The trend of LAD changes in different cell types were plotted using R package ggplot2.

### Epigenomic data analyses

Paired-end fastq data was first processed through a quality control assessment using the FastQC software package (v0.11.9) to evaluate the sequencing quality. The high-quality reads were aligned to the human reference genome hg38 utilizing Bowtie2 (v2.3.5.1), a widely used aligner known for its speed and accuracy. Post-alignment reads with a Mapping Quality score of less than 30 were filtered out using samtools (v1.12) and only non-duplicated mapped reads were retained. For the detection of enriched narrow peaks, indicating regions of significant read accumulation, the MACS2 (Model-based Analysis of ChIP-Seq) (v2.1.2) was employed with its default parameters with IgG as a control. For the detection of enriched broad peaks, the MACS2 (Model-based Analysis of ChIP-Seq) (v2.1.2) was employed with IgG as a control using option “– broad −q 0.05” in addition to default parameters. Broad peaks within 5,000 bp were merged for the downstream analysis. The identified peaks were annotated based on TSS using Homer (v4.11). All intersection calculations were done using bedtools (v2.27.1) software.

### RNA-seq

Total RNA was extracted from RG, IPC and EN nuclei using Quick-RNA FFPE RNA extraction kit (Zymo Research, R1008). Total RNA from NPCs was extracted using Direct-Zol RNA Miniprep kit (Zymo Research, R2052). Samples were depleted of ribosomal RNA (rRNA) using the NEBNext rRNA Depletion kit v2 (NEB, E7400L). Libraries for sequencing were prepared using the NEBNext Ultra II directional library kit (NEB, E7760) following the manufacturer’s protocol. Libraries were quantified by Tapestation D1000 (Agilent) and Qubit high sensitivity dsDNA (Invitrogen) assays, pooled, and sequenced on NovaSeq 6000 or NovaSeq X Plus systems (Illumina).

### Gene expression analyses

All fastq files were aligned to the reference genome (GENCODE GRCh38) by STAR 2.7.11b after adaptor trimming. Reads that passed QC and with read length greater than 50 bp were carried over for downstream counting after mapping to a gene transfer format (GTF) file. Read counts were calculated using featureCounts ^49^. All differential expression analysis was performed by DESeq2 (v1.4) using raw counts across all conditions or cell types. Genes with read count > 50 in at least one sample of a comparison were kept for further analysis. A given gene was considered significantly changing if the false-discovery-rate (FDR) is < 0.05, P value is < 0.05, and Log1p fold-change is ≥ 0.05. Transcripts Per Million (TPM) were calculated to estimate the absolute expression level for each gene. RNA-seq data obtained in this study from GW20 RG, IPC and EN and GW17 RG, IPC and EN from published datasets ^50^ were integrated together into the analysis.

### Gene ontology analyses

For a given list of gene of interest, gene ontology (GO) analysis was performed using clusterProfiler (v3.19) with the default parameters. GO terms with FDR less than 0.05 were designated with significant functional enrichment.

### Statistics and data reproducibility

All the GO-CaRT and RNA-seq experiments in the human cortical plate or germinal zone were performed for at least two biological replicates. No statistical test was used to predetermine the sample size used. P values were derived from Wilcoxon’s rank-sum test for comparisons throughout the study. Common domains of LADs/SPADs and common peaks of histone modifications from two biological replicates were used for the analysis.

### DNA-FISH combined with immunocytochemistry and image analyses

DNA-FISH with ICC for LaminB1 and SON was performed as described previously ^10^ with slight modifications. Briefly, the media was removed from the chambered coverglass containing the dissociated cells from the germinal zone and cortical plate and the cells were fixed with Histochoice MB (Sigma) for 30 min at room temperature. Cells were washed three times with D-PBS, each wash lasting for 10 min. Cells were permeabilized and blocked in D-PBS containing 5% NGS and 0.5% TritonX-100 for 1 h at room temperature. Cells were incubated in antibody binding solution (5% NGS and 0.1% Triton X-100 in D-PBS) containing LaminB1 (Abcam) or SON (Atlas Antibodies) antibodies at 1:500 dilution. The next day, cells were washed three times with D-PBS and incubated with a secondary antibody (Alexa Fluor 647, anti-rabbit, 1:500 dilution) for 2 hours at room temperature. Following three washes with D-PBS, cells were post-fixed in Histochoice MB for 20 min at room temperature. Cells were washed two times with D-PBS, and 0.7% Triton X-100 was included in the third wash to permeabilize the cells. Following a quick wash with D-PBS, cells were treated with a series of ethanol gradient: 70%, 85% and 100% ethanol each for 2 min at room temperature. The chambered coverglass was allowed to dry at 45 °C until all ethanol evaporated. A humidified chamber was prepared using 50% formamide/2X SSC solution in a dark box and pre-heated to 83 °C. The DNA-FISH probe (Empire Genomics) mixture (3-µl probe + 17-µl hybridization buffer + 5-µl denaturation buffer: 70% Formamide, 2X SSC, pH 7.0-8.0) was added to the cells and the chambered coverglass was placed in the pre-heated humidified chamber and denatured at 83 °C for 10 min. Samples were transferred to a 37 °C oven and incubated for 16–24. The next day, wash solution 1 (0.3% NP40 and 0.4X SSC) pre-warmed to 73 °C was added and incubated for 3 min at room temperature. A second wash was performed with wash solution 2 (0.1% NP40 and 2X SSC) for 3 min at room temperature and the coverglass was allowed to dry in dark. Cells were stained with DAPI (1:500 of 1 mg/mL) for 10 min at room temperature, washed with D-PBS and water and mounted with prolong glass (Invitrogen). Samples were left in dark overnight at room temperature to let it cure before imaging.

Imaging was performed using a Leica confocal microscope (Leica TCS SP5X) with 63X oil immersion objective, with a *z*-stack collected for each channel (step size 0.2 μm). ImageJ software (v2.0.0-rc-65/1.52q, build: 961c5f1b7f) was used to process images. Nuclear lamina and speckles were identified by LaminB1 and SON immunostaining, respectively. 3D reconstructions of cells were conducted via Imaris x64 (v9.2.1) software (Bitplane). DNA-FISH dots for probe signals were created at the location of the highest fluorescence intensity via the Spots tool. Spot diameter ranged from 350 nm to 400 nm. Nuclear lamina/speckle surfaces were automatically detected from LaminB1/SON immunostaining with the Surfaces tool. Distance from the center of the FISH spot to the closest nuclear lamina/speckles surface was quantified using the Measurement Points tool. For each experimental condition, 40–60 nuclei were analyzed. In the case that the generated DNA-FISH spot was embedded in the nuclear laminar/speckles surface, distance to the lamina/speckle was quantified as zero.

## Data and code availability

The datasets generated during the current study will be available on public repositories prior to the time of publication. The codes used in the data analyses will be available at GitHub.

## Acknowledgements

This work was supported by NIH grants R01NS112357, R01NS124881, R2NS1125978, VA grant I01BX000252, the Chad Tough Foundation, and the Pathway for Breakthrough in Biomedical Research (PBBR), UCSF. We thank flow cytometry, microscopy, and sequencing cores at UCSF. The UCSF Parnassus Flow CoLab (RRID:SCR_018206) is supported in part by Grant NIH P30 DK063720 and by the NIH S10 Instrumentation Grant S10 1S10OD021822-01. Even Semenza is supported by NSF GRFP fellowship. Li Wang was supported by NIMH grant K99MH131832. The authors thank Tanzila Mukhtar (Arnold Kriegstein lab at UCSF) for her help with primary human tissue samples. Navneet Matharu (UCSF) and members of the Lim laboratory (UCSF) for helpful reading of the manuscript.

## Author Contributions

Conceptualization, S.H.A. and D.A.L.; Methodology, S.H.A., C.Z. E.R.S. and D.A.L.; Software, C.Z.; Formal Analysis, C.Z.; Investigation, S.H.A., C.Z. E.R.S, E.G., M.A.C. and L.W.; Resources, A.R.K. and D.A.L.; Data Curation, C.Z.; Writing – Original Draft, S.H.A., C.Z., and D.A.L.; Writing – Review & Editing, S.H.A., C.Z., E.R.S., E.G., M.A.C., L.W., A.R.K. and D.A.L; Supervision, A.R.K. and D.A.L., Funding Acquisition, A.R.K. and D.A.L.

## Extended Data Figures

**Extended Data Fig. 1.**
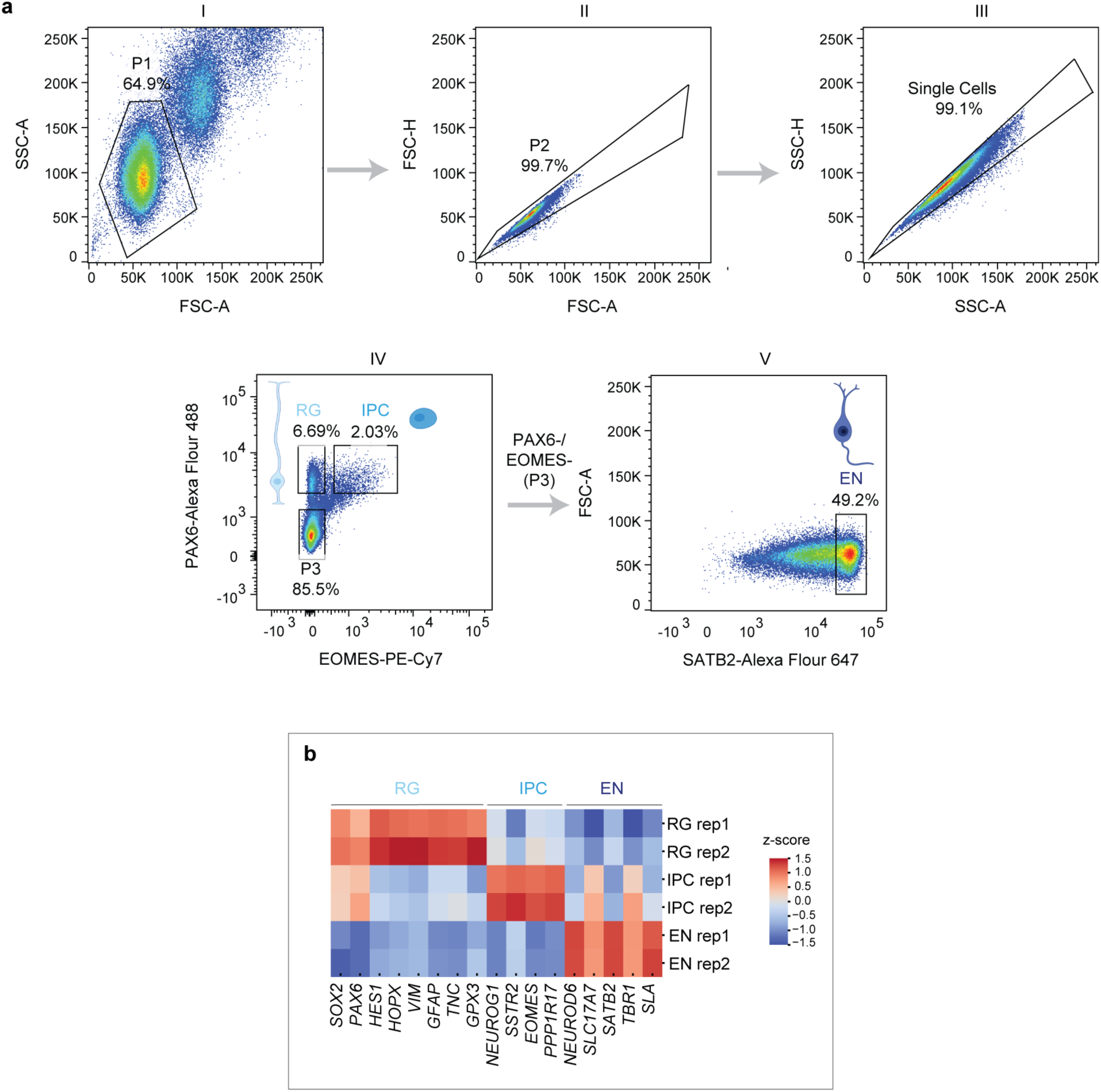
Flow cytometry gating strategy for isolation of RG, IPC and EN and gene expression analysis (related to Figure 1) **a,** Representative gating strategy for FANS-based isolation of RG, IPC and EN from mid-gestational human brain cortex. Nuclei were separated from debris by forward scatter area (FSC-A) versus side scatter area (SSC-A) (P1, plot I). Single cells (singlets) were then selected using two gates based on area versus height of the forward scatter (FSC-A vs. FSC-H, plot II) and side scatter (SSC-A vs. SSC-H, plot III). RG populations were isolated by gating on PAX6-Alexa Fluor 488 staining and IPC by gating on PAX6-Alexa Fluor 488 and EOMES-PE-Cy7 staining (plot IV). EN were isolated from PAX6/EOMES negative population (P3) by gating on SATB2-Alexa Fluor 647 staining (plot V). **b,** Heatmap showing the expression (row normalized) of key marker genes in RG, IPC and EN isolated by FANS from GW20 cortex. R1 and R2 indicate two replicates.

**Extended Data Fig. 2.**
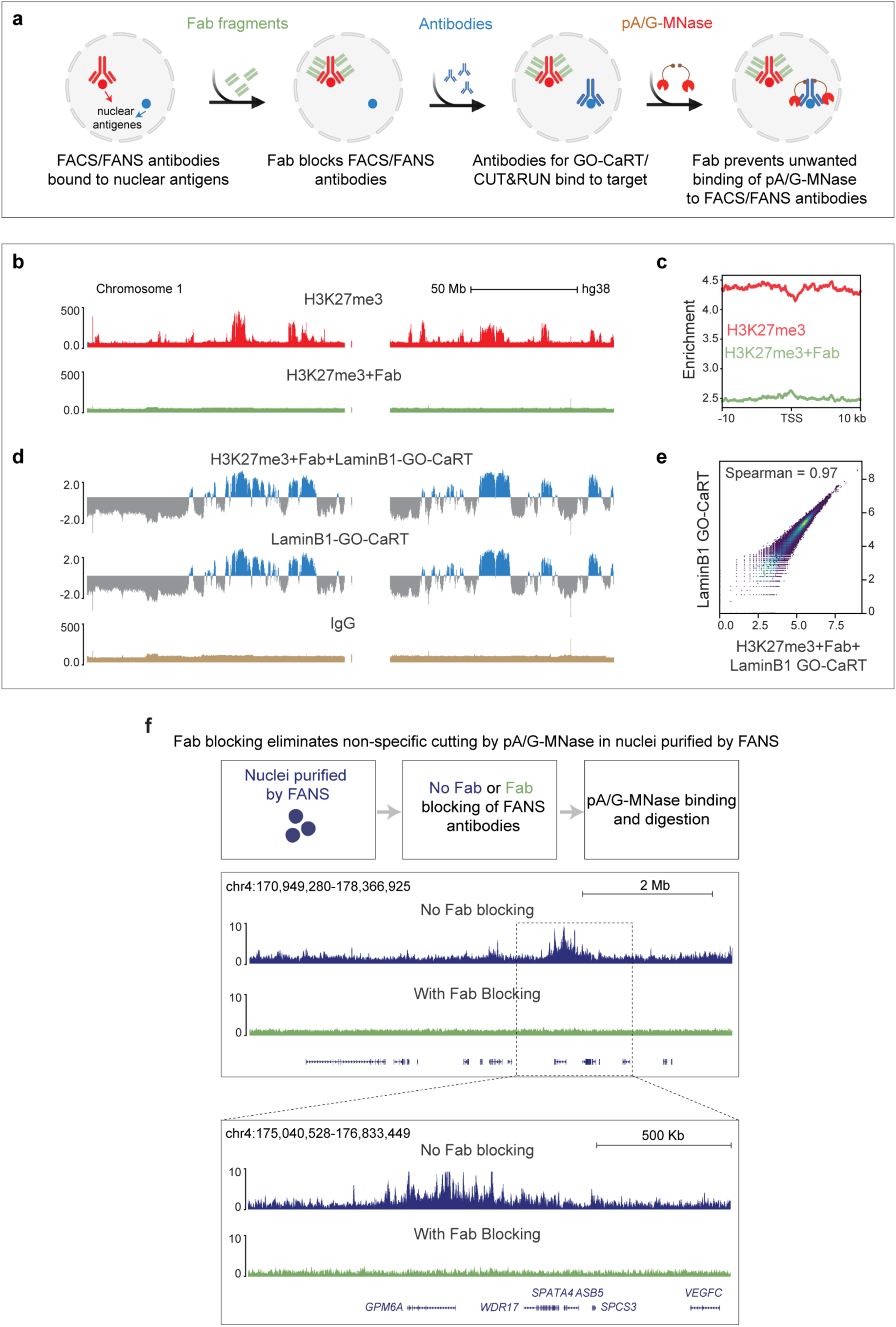
Development of Fab blocking for antibody-based chromatin profiling (related to Figure 1) **a,** Schematic of Fab blocking method. Antibodies used in FANS are blocked by monovalent Fab fragments. In subsequent steps, samples are incubated with antibodies of interest for GO-CaRT and CUT&RUN. The use of Fab blocking prevents interactions between pA/G-MNase, and antibodies used in FANS allowing seamless downstream GO-CaRT and CUT&RUN analyses. **b,** Representative genome browser tracks demonstrating the efficacy of Fab fragments for blocking interaction between pA/G-MNase and antibodies to H3K27me3 in CUT&RUN assay in HEK293T cells. Without Fab treatment, CUT&RUN reveals expected levels of H3K27me3 enrichment (red track). In cells treated with Fab fragments, the H3K27me3 signal is lost genome-wide (green track). **c,** Genome wide enrichment of H3K27me3 with or without Fab blocking. **d,** Fab blocking does not interfere with subsequent GO-CaRT with LaminB1 antibody. Top blue track: HEK293T cells in which Fab fragments were used to block H3K27me3 antibodies and subsequently incubated with antibodies to LaminB1 protein for LaminB1 GO-CaRT analysis. Bottom blue: HEK293T cells that were directly subjected to LaminB1 GO-CaRT analysis. **e,** Genome-wide scatter plot showing Spearman correlation between LaminB1 GO-CaRT in HEK293T cells after Fab blocking of H3K27me3 vs. those that were directly subjected to LaminB1 GO-CaRT. **f,** Fab blocking eliminates the background signal produced by non-specific targeting of pA/G-MNase by antibodies used in FANS. Representative tracks generated by pA/G-MNase digestion with or without Fab blocking of FANS-isolated EN.

**Extended Data Fig. 3.**
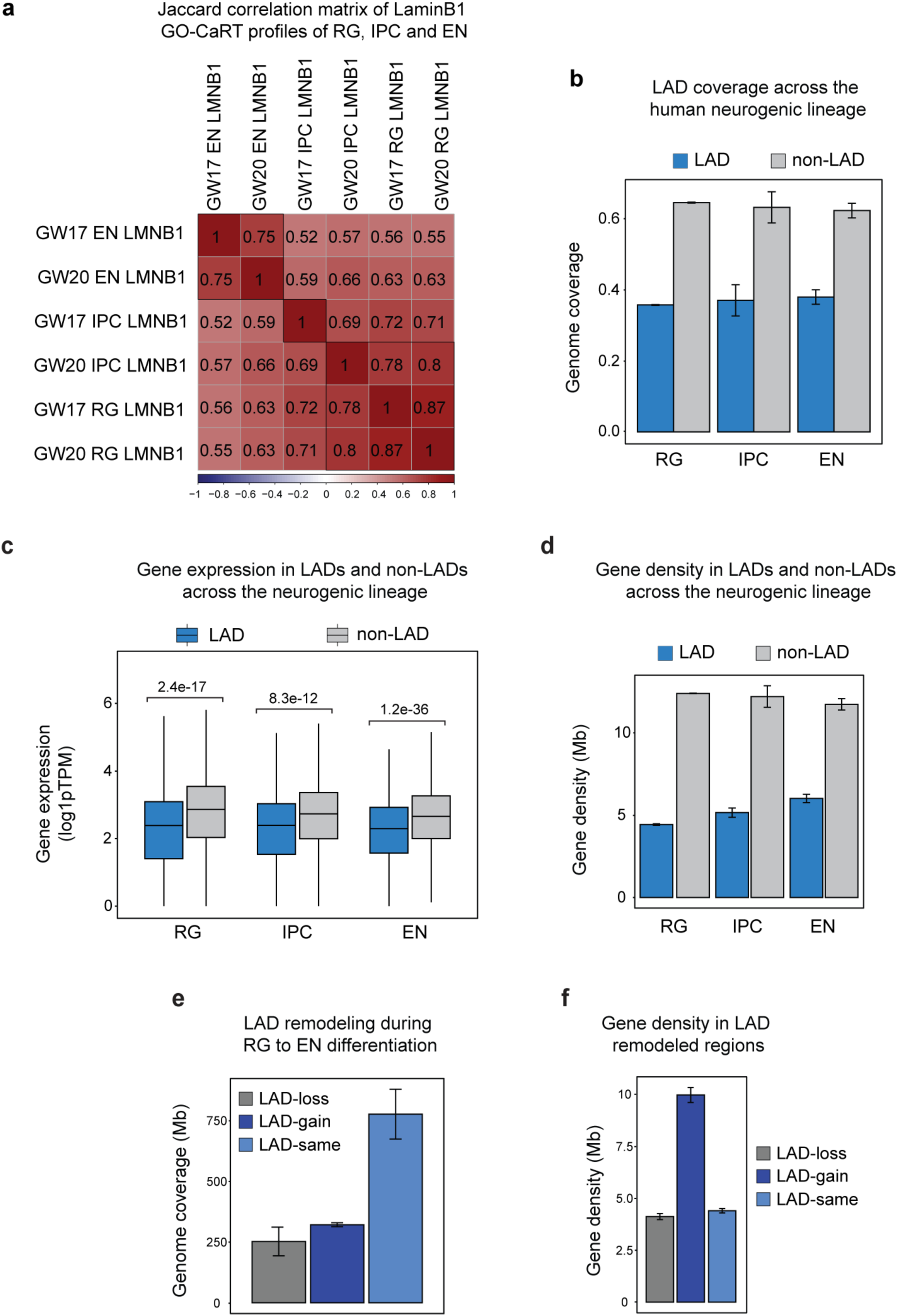
Various genomic features of LADs across the neurogenic lineage (related to Figure 1) **a,** Jaccard correlation between LaminB1 GO-CaRT of RG, IPC and EN isolated by FANS from GW17 and GW20 brain cortex. **b,** Percent LAD coverage in RG, IPC and EN. Standard deviation between GW17 and GW20 time points. **c,** Box plot depicting average gene expression in LADs (RG, n= 4947; IPC, n= 5462; EN, n= 6929) and non-LADs (RG, n= 24593; IPC, n= 24057; EN, n= 22590) as determined by RNA-seq. *n*, number of genes. *P* values determined by Wilcoxon rank sum test. Boxes show the range from lower (25th percentile) to upper quartiles (75th percentile), with the median line (50th percentile); whiskers extend 1.5 times the inter-quartile range from the bounds of the box. **d,** Gene density in LADs and non-LADs across the neurogenic lineage. **e,** Genome coverage of LAD remodeling during the differentiation of RG to EN. **f,** Gene density in LADs that are lost, gained, or remain unchanged during RG to EN differentiation.

**Extended Data Fig. 4.**
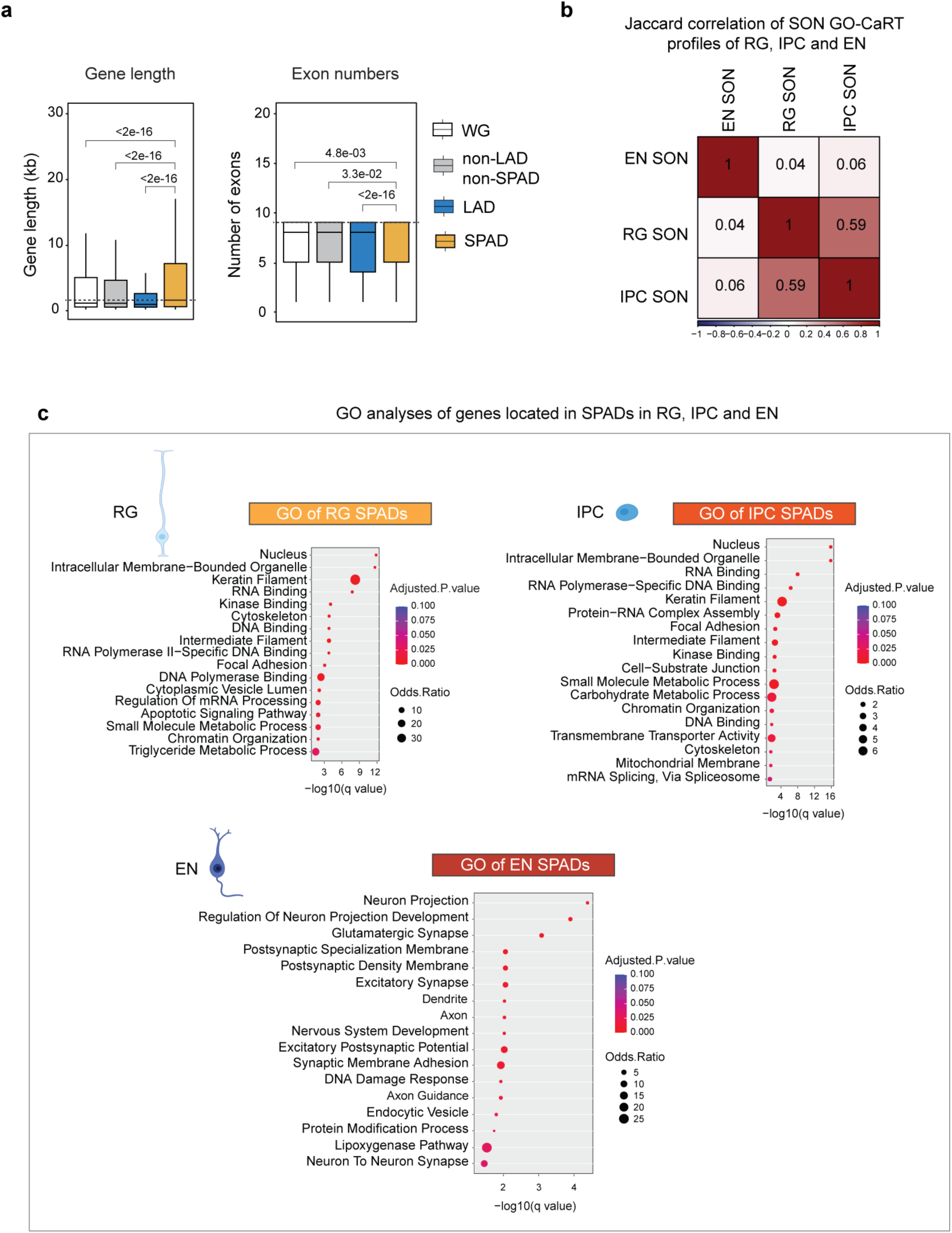
Analysis of LAD to SPAD remodeling during RG to EN differentiation (related to Figure 2) **a,** Box plot showing average gene length and exon number across the neurogenic lineage in different genomic compartments (LADs, non-LAD/non-SPADs and SPADs) as compared to the whole genome. *P* values determined by Wilcoxon rank sum test. Boxes show the range from lower quartiles (25th percentile) to upper quartiles (75th percentile), with the median line (50th percentile); whiskers extend 1.5 times the interquartile range from the bounds of the box. **b,** Jaccard correlation matrix of SON GO-CaRT profiles of RG, IPC and EN at GW17. **c,** GO analyses of RG, IPC and EN SPADs. GO terms were sorted based on their significance (−log10(*q* value)); the size of the bubble represents the Odds Ratio for each term.

**Extended Data Fig. 5.**
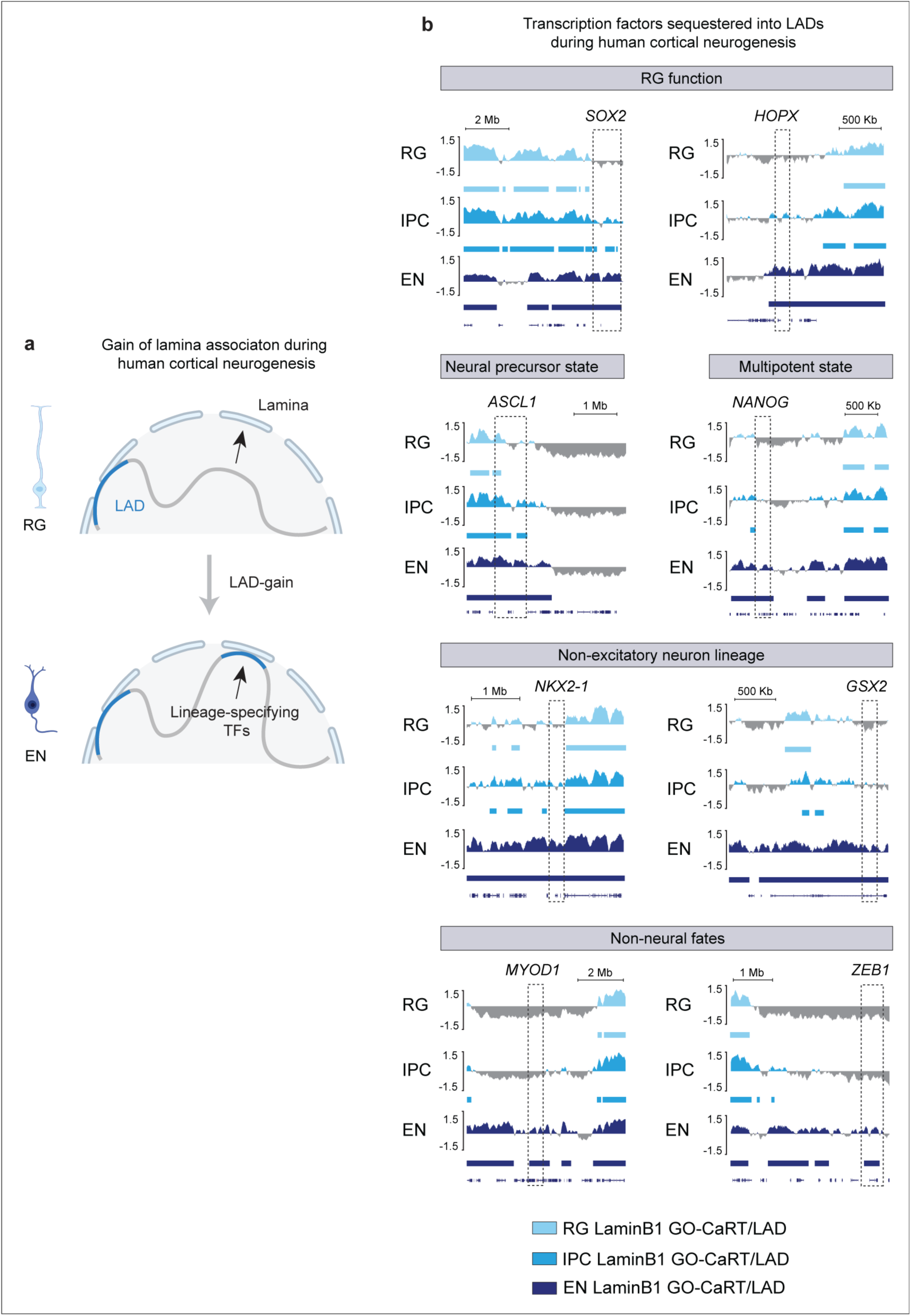
Genes important for various lineages are sequestered into LADs during cortical neurogenesis (related to Figure 3) **a,** A cartoon illustrating the gain of lamina association (LAD-gain) during cortical neurogenesis. TF, transcription factors **b,** LaminB1 GO-CaRT tracks in RG, IPC and EN at genes that are sequestered into LADs during cortical neurogenesis. LADs are shown below the track by rectangles. Dashed boxes show indicated gene loci.

**Extended Data Fig. 6.**
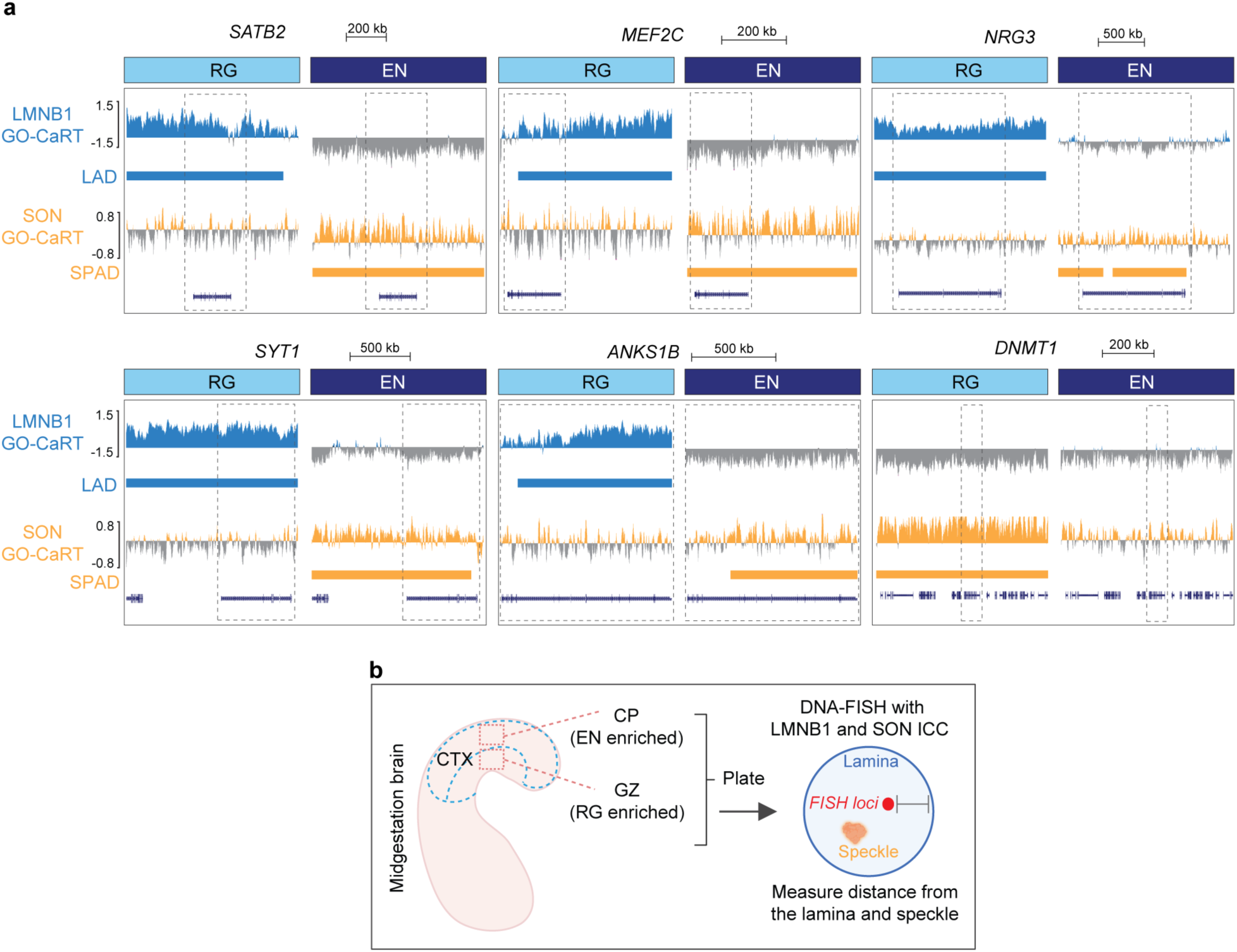
DNA-FISH analyses of genes that move from lamina to speckles during cortical neurogenesis (related to Figure 3) **a,** LaminB1 and SON GO-CaRT tracks in RG and EN over gene loci analyzed by DNA-FISH. **b,** Cartoon showing the experimental design for DNA-FISH.

**Extended Data Fig. 7.**
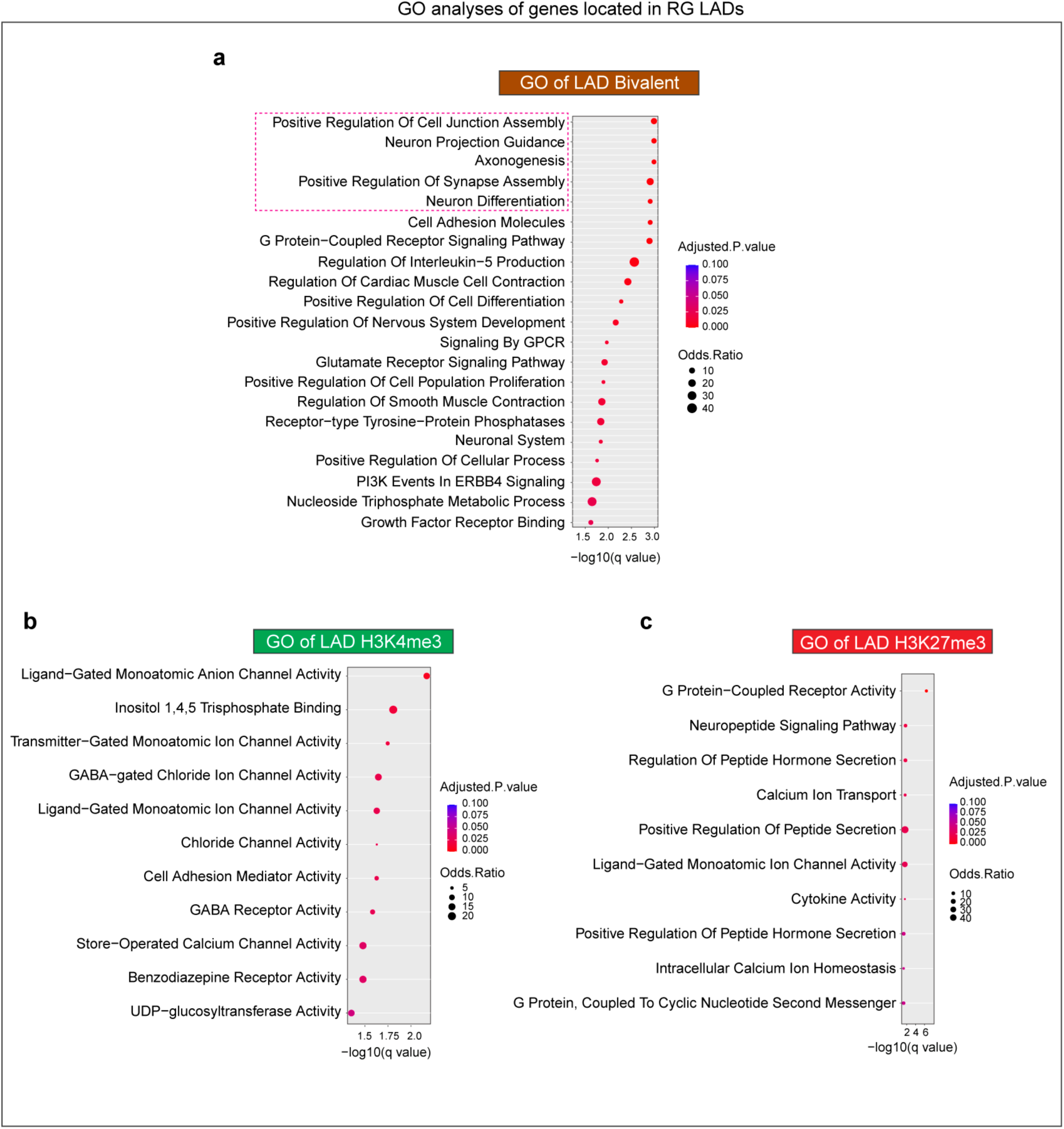
RG LADs are enriched in neurodevelopmental genes marked by bivalent chromatin (related to Figure 4) **a,** GO analyses of bivalent genes in RG LADs. **b,** GO analyses of H3K4me3-monovalent genes in RG LADs. **(C)** GO analyses of H3K27me3-monovalent genes in RG LADs. GO terms were sorted based on their significance (−log10(*q* value)); the size of the bubble represents the Odds Ratio for each term.

**Extended Data Fig. 8.**
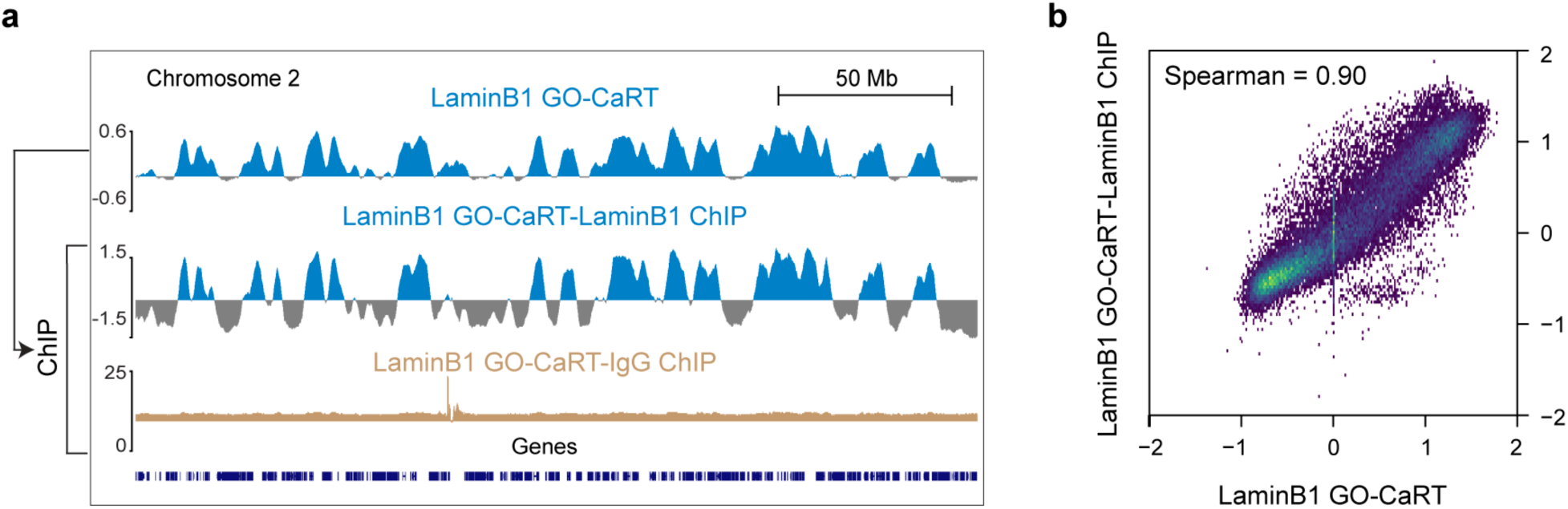
LaminB1 GO-CaRT.ChIP-reChIP in iPSC-derived NPCs (related to Figure 5) **a,** Representative genome tracks of LaminB1 GO-CaRT and subsequent LaminB1 ChIP and IgG ChIP performed on LaminB1 GO-CaRT supernatant (input). **b,** Genome-wide Spearman correlation between LaminB1 GO-CaRT input and LaminB1 GO-CaRT-LaminB1 ChIP.

**Extended Data Fig. 9.**
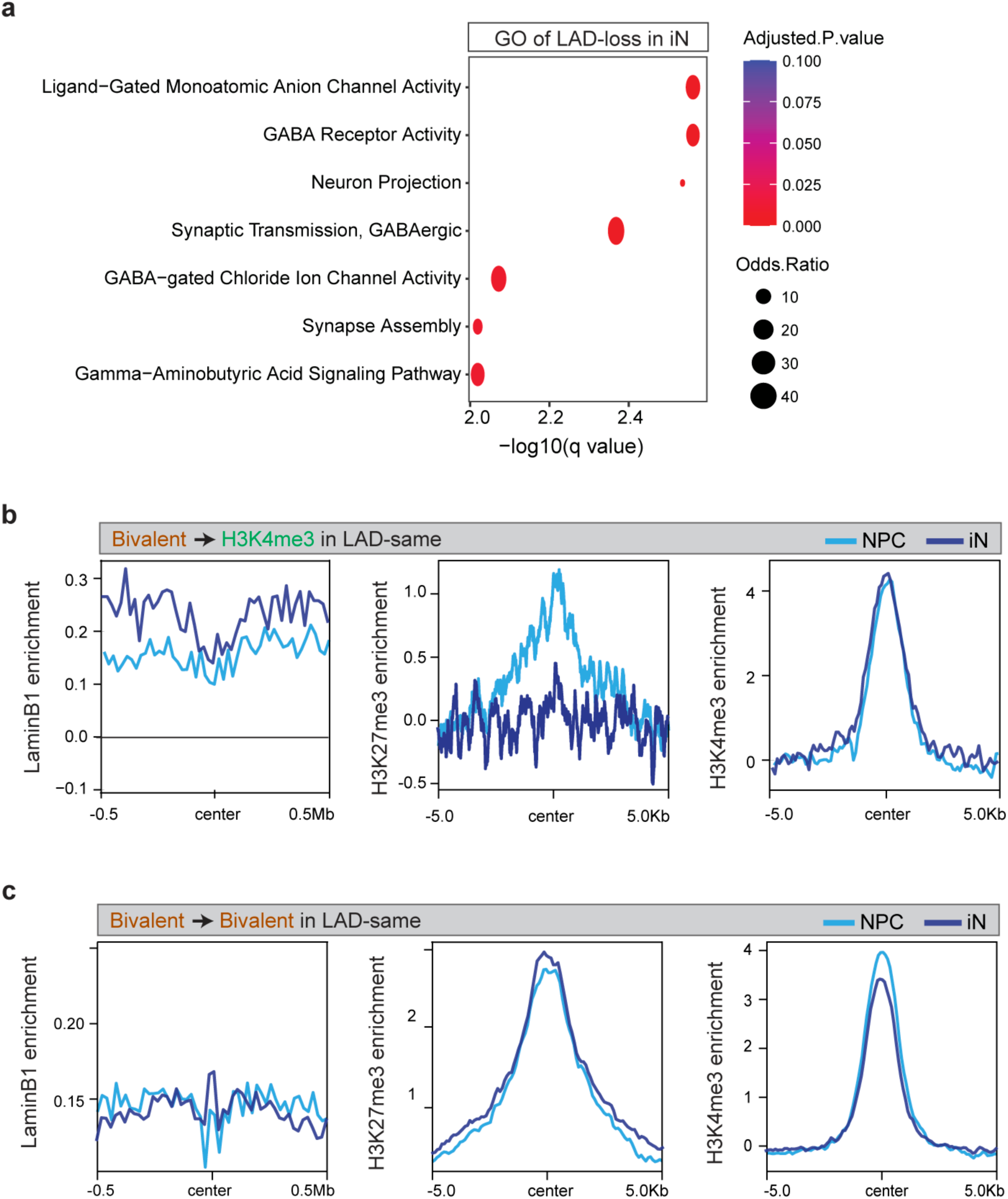
LAD dynamics and regulation of bivalent chromatin during *in vitro* neurogenesis (related to Figure 6) **a,** GO analyses of genes that reside in genomic regions that undergo LAD-loss during *in vitro* neurogenesis from (during NPC to iN differentiation). GO terms were sorted based on their significance (−log10(*q* value)); the size of the bubble represents the Odds Ratio for each term. **B,** Normalized LaminB1, H3K27me3 and H3K4me3 levels for bivalent genes that resolve to the H3K4me3-monovalent state in LAD-same regions during NPC to iN differentiation. **c,** Normalized LaminB1, H3K27me3 and H3K4me3 levels for bivalent genes that retain bivalency in LAD-same regions during NPC to iN differentiation.

**Extended Data Fig. 10.**
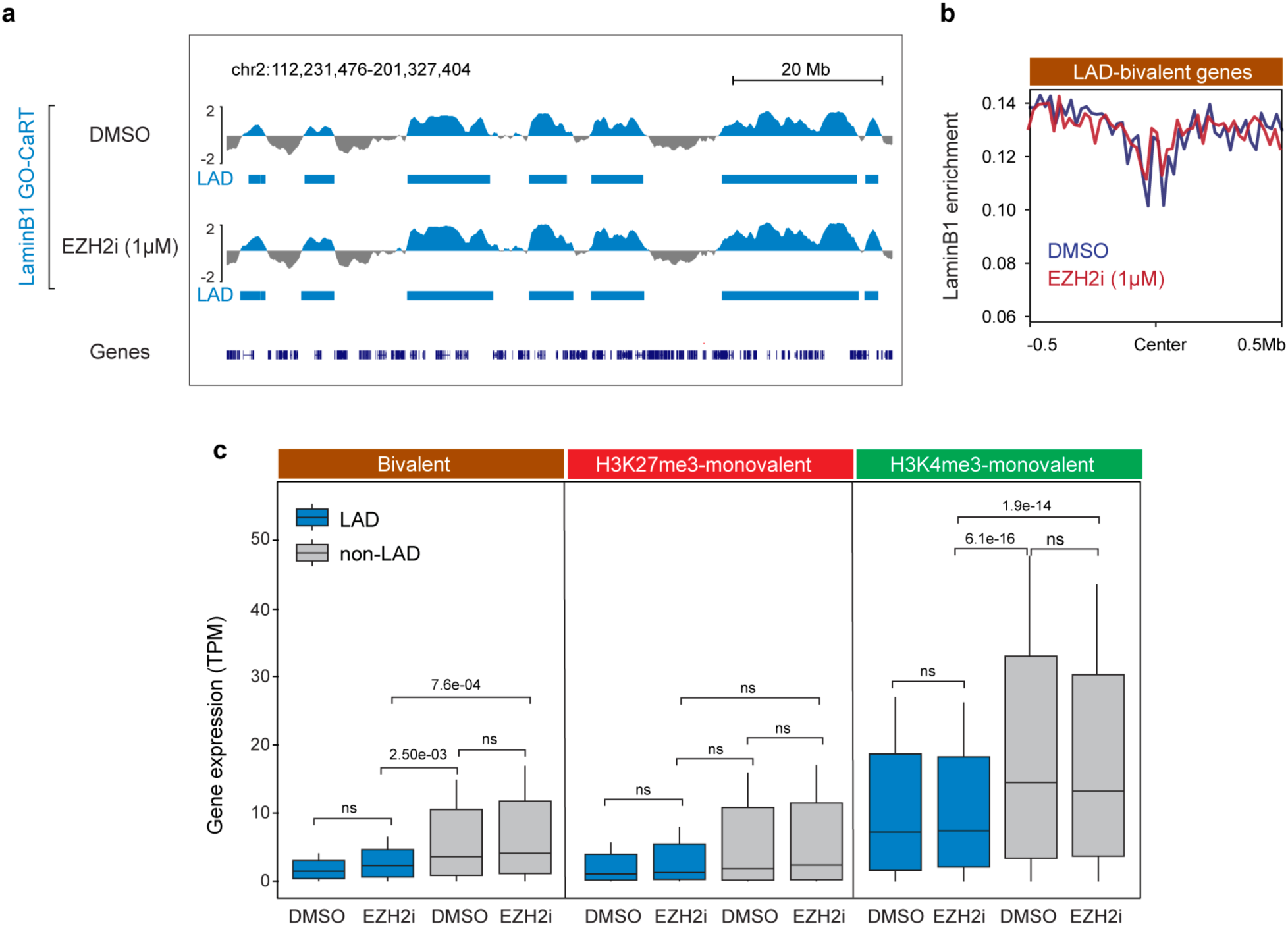
EZH2 inhibition in NPCs does not grossly alter the LAD architecture and expression of LAD resident genes (related to Figure 7) **a,** Representative LaminB1 GO-CaRT tracks in NPCs treated with DMSO or 1 μM Tazemetostat (EZH2i) for 7 DIV. LADs are shown below the track **b,** Normalized LaminB1 levels at LAD bivalent genes in NPCs treated with DMSO and 1 μM Tazemetostat (EZH2i) for 7 DIV. **c,** Box plot showing average gene expression of bivalent (n = 2158), H3K27me3-monovalent (n = 1476) and H3K4me3-monovalent (n = 9550) genes in LADs and non-LADs in NPCs treated with DMSO or 1 μM Tazemetostat (EZH2i) for 7 DIV. *n*, number of genes across the whole genome; *P* values determined by Wilcoxon rank sum test. Boxes show the range from lower quartiles (25th percentile) to upper quartiles (75th percentile), with the median line (50th percentile); whiskers extend 1.5 times the interquartile range from the bounds of the box.

**Extended Data Table 1.** Reagents and resource information.

| Reagent or Resource | Source | Identifier |
| --- | --- | --- |
| **Antibodies** |  |  |
| Alexa Fluor® 488 Mouse anti-Human Pax-6 | BD Biosciences | 561664 |
| EOMES Monoclonal Antibody (WD1928), PE-Cyanine7 | Invitrogen | 25-4877-42 |
| Recombinant Alexa Fluor® 647 Anti-SATB2 antibody | Abcam | ab196536 |
| AffiniPure Fab Fragment Donkey Anti-Rabbit IgG (H+L) | Jackson Immuno | 711-007-003 |
| AffiniPure Fab Fragment Donkey Anti-Mouse IgG (H+L) | Jackson Immuno | 715-007-003 |
| Anti-Lamin B1 antibody | Abcam | ab16048 |
| Anti-SON antibody | Atlas antibodies | HPA031755 |
| Tri-Methyl-Histone H3 (Lys27) (C36B11) | Cell Signaling Technology | 9733 |
| Tri-Methyl-Histone H3 (Lys4) (C42D8) Rabbit | Cell Signaling Technology | 9751 |
| Normal Rabbit IgG | Cell Signaling Technology | 2729S |
| Guinea Pig anti-Rabbit IgG | Antibodies-online.com | ABIN101961 |
| **Peptides, recombinant proteins, chemicals** |  |  |
| Mouse Lamin B1 peptide | Abcam | ab16375 |
| Histone H3K27me3 Trimethyl Peptide | EpigenTek | R-1034-200 |
| Histone H3K4me3 Trimethyl Peptide | EpigenTek | R-1022-200 |
| pA/G-MNase | Macro Lab (UC Berkley) | - |
| pA-Tn5 | Macro Lab (UC Berkley) | - |
| Proteinase K Solution (20 mg/mL) | Thermofisher | AM2548 |
| DNase Free RNase A | Thermofisher | EN0531 |
| RiboLock RNase Inhibitor | Thermofisher | EO0382 |
| RNasin® Plus Ribonuclease Inhibitor | Promega | N2615 |
| Glycogen, RNA grade | Thermofisher | R0551 |
| Spermidine >99% (GC) | Sigma | S2626-1G |
| ProLong™ Glass Antifade Mountant | Thermofisher | P36980 |
| Tazemetostat (EPZ-6438) | Selleckchem | S7128 |
| UltraPure™ Phenol:Chloroform:Isoamyl Alcohol (25:24:1, v/v) | Thermofisher | 15593031 |
| Pierce™ 16% Formaldehyde (w/v), Methanol-free | Thermofisher | 28906 |
| HistoChoice® Tissue Fixative | Millipore Sigma | H2904 |
| Halt™ Protease Inhibitor Cocktail (100X) | Thermofisher | 78438 |
| Digitonin solution- high purity | Millipore Sigma | CHR103 |
| Trizol reagent | Thermofisher | 15596018 |
| Dynabeads® Protein A | Thermofisher | 10002D |
| BioMag®Plus Concanavalin A | Polysciences | 86057-10 |
| RNAClean XP | Beckman Coulter | A63987 |
| Agencourt AMPure XP | Beckman Coulter | A63881 |
| SPRIselect Reagent<br>ReLeSR | Beckman Coulter<br>StemCell Technologies | B23318<br>5872 |
| StemPro Accutase | Thermofisher | A1110501 |
| Matrigel® Growth Factor Reduced (GFR)<br>Basement Membrane Matrix | Corning | 356231 |
| Laminin | Millipore Sigma | L2020-1MG |
| Fibronectin bovine plasma | Millipore Sigma | F4759-1MG |
| Poly-L-ornithine hydrobromide | Millipore Sigma | P3655-50MG |
| Poly-D-lysine hydrobromide | Millipore Sigma | P0899-50MG |
| StemFlex Medium | Thermofisher | A3349401 |
| DMEM/F-12, GlutaMAX™ supplement | Thermofisher | 10565042 |
| Neurobasal-A Medium | Thermofisher | 10888022 |
| B27 minus Vitamin A | Invitrogen | 12587-010 |
| N2 supplement | Life Technologies | 17502-048 |
| DMEM | Corning | 10-017-CV |
| FBS | VWR | 97068-085 |
| Antibiotic/antimycotic 100X | Thermofisher | 15240062 |
| L-Glutamine 200mM | Thermofisher | 25030081 |
| Y-27632 ROCK inhibitor | Selleckchem | S1049 |
| Recombinant Human EGF | PeproTech | AF-100-15 |
| Recombinant Human FGF-basic | PeproTech | 100-18B |
| Insulin-Transferrin-Selenium (ITS-G) | Thermofisher | 41400045 |
| L-Ascorbic acid 2-phosphate<br>sesquimagnesium salt | Santa Cruz Biotech | sc-228390 |
| LDN-193189 | Selleckchem | S2618 |
| SB431542 | Selleckchem | S1067 |
| XAV939 | Cayman Chemical | 13596 |
| N6,2'-O-Dibutyryl adenosine 3',5'-cyclic<br>monophosphate sodium salt | MilliporeSigma | D0627-100MG |
| Recombinant Human/Murine/Rat BDNF | PeproTech | 450-02 |
| Recombinant Human GDNF | PeproTech | 450-10 |
| DAPT | MedChemExpress | HY-13027 |
| **Critical commercial assays** |  |  |
| KAPA HyperPrep Kit with KAPA Library<br>Amplification Primer Mix (10X) | Kapa | KK8504 |
| TruSeq DNA Single Indexes Set A (12<br>Indexes, 24 Samples) | Illumina | 20015960 |
| TruSeq DNA Single Indexes Set B (12<br>Indexes, 24 Sample) | Illumina | 20015961 |
| NEBNext® rRNA Depletion Kit v2<br>(Human/Mouse/Rat) | New England Biolabs | E7400L |
| NEBNext® Ultra™ II Directional RNA Library Prep with Sample Purification Beads | New England Biolabs | E7765S |
| NEBNext® Multiplex Oligos for Illumina® (Index Primers Set 3) | New England Biolabs | E7710S |
| NEBNext® Multiplex Oligos for Illumina® (Index Primers Set 4) | New England Biolabs | E7730S |
| Papain Dissociation System | Worthington | LK003150 |
| StemXVivo Neural Progenitor Differentiation Kit | R&D systems | SC035 |
| AbC™ Total Antibody Compensation Bead Kit | Thermofisher | A10513 |
| Qubit dsDNA HS Assay Kit | Thermofisher | Q32854 |
| RNA ScreenTape | Agilent | 5067-5576 |
| RNA ScreenTape Sample Buffer | Agilent | 5067-5577 |
| D1000 ScreenTape | Agilent | 5067-5582 |
| D1000 Reagents | Agilent | 5067-5583 |
| High Sensitivity D1000 ScreenTape | Agilent | 5067-5584 |
| High Sensitivity D1000 Reagents | Agilent | 5067-5585 |
| Direct-Zol RNA Miniprep | Zymo research | R2052 |
| Quick-RNA FFPE Miniprep | Zymo research | R1008 |
| Phase Lock Gel Heavy | VWR | 10847-802 |
| 8-well Chambered Coverglass w/ non-removable wells (1.5 Borosilicate Glass) | Thermofisher | 155409PK |
| **Experimental models: Cell lines** |  |  |
| HEK293T | ATCC |  |
| WTC11-hiPSC | Conklin lab; Kreitzer et al., 2013 |  |
| H1-ESC | WiCell (authenticated at source) |  |
| **DNA-FISH probes (genomic co-ordinates, hg38)** |  |  |
| NRG3 locus (chr10:81994239-82200685) | Empire Genomics | RP11-828D16 |
| SYT1 locus (chr12:79120096-79278995) | Empire Genomics | RP11-76L7 |
| ANKS1B locus (chr12:99144442-99346123) | Empire Genomics | RP11-315N20 |
| SATB2 locus (chr2:199280574-199450448) | Empire Genomics | RP11-755G8 |
| MEF2C locus (chr5:88738836-88895936) | Empire Genomics | RP11-1147F22 |
| DNMT1 locus (chr19:10027189-10224063) | Empire Genomics | RP11-298C17 |
| **Plasmids** |  |  |
| pAG/MNase | Addgene | 123461 |
| 3XFlag-pA-Tn5-Fl | Addgene | 124601 |

